# Combinatorial immune refocusing within the influenza hemagglutinin head elicits cross-neutralizing antibody responses

**DOI:** 10.1101/2023.05.23.541996

**Authors:** Annie Dosey, Daniel Ellis, Seyhan Boyoglu-Barnum, Hubza Syeda, Mason Saunders, Michael Watson, John C. Kraft, Minh N. Pham, Miklos Guttman, Kelly K. Lee, Masaru Kanekiyo, Neil P. King

**Author notes:** Correspondence (N.P.K.).

## Abstract

The head domain of influenza hemagglutinin (HA) elicits potently neutralizing yet mostly strain-specific antibodies during infection and vaccination. Here we evaluated a series of immunogens that combined several immunofocusing techniques for their ability to enhance the functional breadth of vaccine-elicited immune responses. We designed a series of “trihead” nanoparticle immunogens that display native-like closed trimeric heads from the HAs of several H1N1 influenza viruses, including hyperglycosylated variants and hypervariable variants that incorporate natural and designed sequence diversity at key positions in the periphery of the receptor binding site (RBS). Nanoparticle immunogens displaying triheads or hyperglycosylated triheads elicited higher HAI and neutralizing activity against vaccine-matched and -mismatched H1 viruses than corresponding immunogens lacking either trimer-stabilizing mutations or hyperglycosylation, indicating that both of these engineering strategies contributed to improved immunogenicity. By contrast, mosaic nanoparticle display and antigen hypervariation did not significantly alter the magnitude or breadth of vaccine-elicited antibodies. Serum competition assays and electron microscopy polyclonal epitope mapping revealed that the trihead immunogens, especially when hyperglycosylated, elicited a high proportion of antibodies targeting the RBS, as well as cross-reactive antibodies targeting a conserved epitope on the side of the head. Our results yield important insights into antibody responses against the HA head and the ability of several structure-based immunofocusing techniques to influence vaccine-elicited antibody responses.

**HIGHLIGHTS:**

- Generalization of trihead antigen platform to several H1 hemagglutinins, including hyperglycosylated and hypervariable variants
- Trimer-stabilizing mutations in trihead nanoparticle immunogens lead to lower levels of non-neutralizing antibody responses in both mice and rabbits
- Hyperglycosylated triheads elicit higher antibody responses against broadly neutralizing epitopes

## INTRODUCTION

Influenza viruses currently persist as a major public health threat due to their high evolutionary rate that gives rise to frequent antigenic drift amongst circulating strains (Carrat and Flahault 2007; Bedford et al. 2014). This strain divergence is due in large part to diversity in the surface glycoprotein HA as a result of immune pressure (Koel et al. 2013; Rejmanek et al. 2015). Despite its variation amongst strains, HA characterization has revealed two functionally conserved sites: the RBS in the head domain, which mediates host cell entry by binding to sialic acid on host glycoproteins, and a conserved antigenic region in the HA stem that is involved in host membrane fusion (Wu and Wilson 2017). The identification of several bNAbs against these sites has made them central targets in vaccine design efforts (Ekiert et al. 2012; Whittle et al. 2011; P. S. Lee et al. 2012; Krause et al. 2011; Dreyfus et al. 2012; Kallewaard et al. 2016; Li et al. 2022; McCarthy et al. 2018). The HA head is immunodominant and antibodies that bind near the RBS typically exhibit potent neutralization by blocking receptor binding (Altman et al. 2015; Wu and Wilson 2017; Angeletti et al. 2017). However, high levels of antigenic variation in the head domain allow influenza viruses to evade head-directed immunity through antigenic drift (Altman, Angeletti, and Yewdell 2018). Thus, bNAbs targeting the conserved RBS itself are rarely elicited by infection or vaccination, and response breadth is often limited by frequent residue mutations in the RBS periphery (Zost et al. 2019). By contrast, antibodies targeting the central stem epitope tend to react more broadly to HAs from different influenza viruses, but these antibodies are not always neutralizing and they are difficult to elicit due to the immune subdominant nature of the stem. Recently, broadly protective antibodies against additional antigenic sites on HA have been discovered, including the stem anchor epitope (Guthmiller et al. 2022; Benton et al. 2018) and the trimer interface in the head domain (Watanabe et al. 2019; Bangaru et al. 2019; J. Lee et al. 2016). The broad binding of trimer interface-directed antibodies against divergent HA subtypes makes this site an intriguing vaccine target. Although they lack neutralizing activity *in vitro*, initial reports indicate that trimer interface-directed antibodies can be protective in animal models (Watanabe et al. 2019; Bangaru et al. 2019).

In addition to amino acid substitutions, influenza viruses use glycosylation of their surface proteins as a mechanism of immune evasion. Analysis of HA sequences obtained over the past century has revealed that the HA head has acquired more glycans over time, and this has been shown to both prevent antibody binding and to lower the number of mutations required for immune evasion (Kobayashi and Suzuki 2012; Wei et al. 2010). These observations have inspired the use of glycan engineering in vaccine design as a means to direct immune responses to target epitopes. Previous studies have generated hyperglycosylated immunogens by introducing new N-linked glycosylation motifs in the immunodominant head domain, resulting in the diversion of antibody responses onto conserved, subdominant epitopes in either the stem domain (Eggink, Goff, and Palese 2014) or the trimer interface (Bajic et al. 2019). A reduction in non-neutralizing trimer interface responses was seen in a separate study that combined hyperglycosylation and disulfide bond engineering in full-length HA ectodomains, but this did not increase the elicitation of broadly-reactive RBS-directed antibodies (Thornlow et al. 2021).

The utilization of variable sequences within vaccines as a means to focus immune responses onto conserved epitopes has been implemented in various formats. One approach threaded divergent HA sequences onto H3 or B HA antigens and then employed those varying immunogens in a sequential vaccination regimen, which resulted in protection from vaccine-mismatched challenge (Sun Weina et al. 2019; Broecker et al. 2019). Beyond heterologous prime-boost strategies, another strategy is the co-display of several variants of a given antigen on nanoparticle scaffolds, an approach known as “mosaic nanoparticle display.” In the first application of this strategy, the assembly of up to eight different HA RBDs on the same mosaic ferritin nanoparticle resulted in higher levels of cross-reactive B cells and significantly better neutralization breadth against a large panel of H1 viruses compared to the same RBD antigens presented in either a heterologous prime-boost regimen or as a cocktail of homotypic (i.e., single-strain) nanoparticles (Kanekiyo et al. 2019). Furthermore, a recent study applying this mosaic approach to the display of trimeric HA ectodomains elicited broadly protective immune responses that were subtly yet consistently superior to cocktails of homotypic particles (Boyoglu-Barnum et al. 2021), although another study using a different display approach failed to observe a similar effect (Cohen, Yang, et al. 2021). Mechanistically, it has been proposed that B cell receptors (BCRs) targeting conserved epitopes on mosaic nanoparticles have an avidity advantage compared to BCRs directed against variable epitopes, and that this allows for increased antigen binding and thus B cell activation (Kanekiyo et al. 2019). To date, all reports of mosaic nanoparticle immunogens have tested co-display of antigens with wild-type sequences from various virus isolates; the use of synthetic or designed antigenic variation in mosaic nanoparticles remains unexplored.

Here we evaluated the ability of several known and novel immunofocusing techniques—conformational stabilization, hyperglycosylation, mosaic nanoparticle display, and the design of synthetic “hypervariable” antigens—to focus antibody responses on the conserved RBS of the HA head. We found that combining conformational stabilization, hyperglycosylation, and mosaic nanoparticle display substantially altered the epitopes targeted by vaccine-elicited antibodies to elicit potent vaccine-matched and -mismatched responses, while additional co-display of hypervariable antigens did not further alter the potency or epitope specificity of the serum antibody response.

## RESULTS

### Design and Immunogenicity of Hyperglycosylated Trihead Nanoparticle Immunogens

In the accompanying manuscript we describe the design of a “trihead” antigen in which the head domain of H1 A/New Caledonia/20/1999 (NC99) was stabilized in a native-like trimeric state via hydrophobic mutations at the trimer interface and a rigid fusion to the trimeric component of the I53_dn5 nanoparticle (Ellis et al., n.d.). We also show that this stabilized immunogen, which we refer to here as TH-NC99 (**Figure 1A**), elicits potent neutralization and HAI activity in immunized mice. To maximize focus of immune responses elicited by this immunogen to the conserved RBS, three and five additional N-linked glycans were engineered into epitopes distant from the RBS to create the hyperglycosylated NC99 triheads TH-NC99-7gly and TH-NC99-9gly, respectively (**Figure 1A**). The new glycans were designed using the Rosetta modeling suite, both to guide sequon design and model glycan structure (Adolf-Bryfogle et al. 2021; Leman et al. 2020). TH-NC99 constructs with individual glycan additions were first evaluated for their secretion from HEK293F cells (data not shown) and those that maintained expression were then combined. We also generated “monohead” antigens lacking the trimer interface-stabilizing mutations and rigidifying disulfide bond, which form monomers with exposed trimer interfaces (Ellis et al., n.d.), bearing the four wild-type (MH-NC99) and five additional (MH-NC99-9gly) N-linked glycans. Amino acid sequences for all novel proteins used in this study can be found in **Table S1**. The monohead and trihead antigens were secreted from HEK293F cells as genetic fusions to the I53_dn5B trimer and purified via immobilized metal affinity chromatography (IMAC) and size exclusion chromatography (SEC) (**Figure S1A**). SDS-PAGE revealed slower migration for the hyperglycosylated monohead and trihead subunits compared to their wild-type counterparts (**Figures 1B** and **S1B**). Treatment of the trimeric components with PNGaseF resulted in a large decrease in apparent molecular weight by SDS-PAGE and uniform migration of all constructs. Biolayer interferometry (BLI) using the anti-RBS bNAb C05 (Ekiert et al. 2012) showed similar binding profiles for all five components, indicating that the hyperglycosylated antigens maintain RBS antigenicity (**Figure 1C**). By contrast, BLI using the anti-trimer interface mAb FluA-20 (Bangaru et al. 2019) showed high binding to monohead components but minimal binding to all trihead components, confirming stable closure of the trimer interface in the trihead antigens. Lastly, BLI using the anti-lateral patch mAb Ab6649 (Raymond et al. 2018) showed nearly full binding to TH-NC99-7gly but greatly diminished binding to TH-NC99-9gly, likely due to the additional glycan at position 167 in TH-NC99-9gly that is in the center of the Ab6649 epitope. MH-NC99-9gly showed moderate Ab6649 binding, possibly due to less consistent glycosylation at that position than in TH-NC99-9gly.

**Figure 1.**
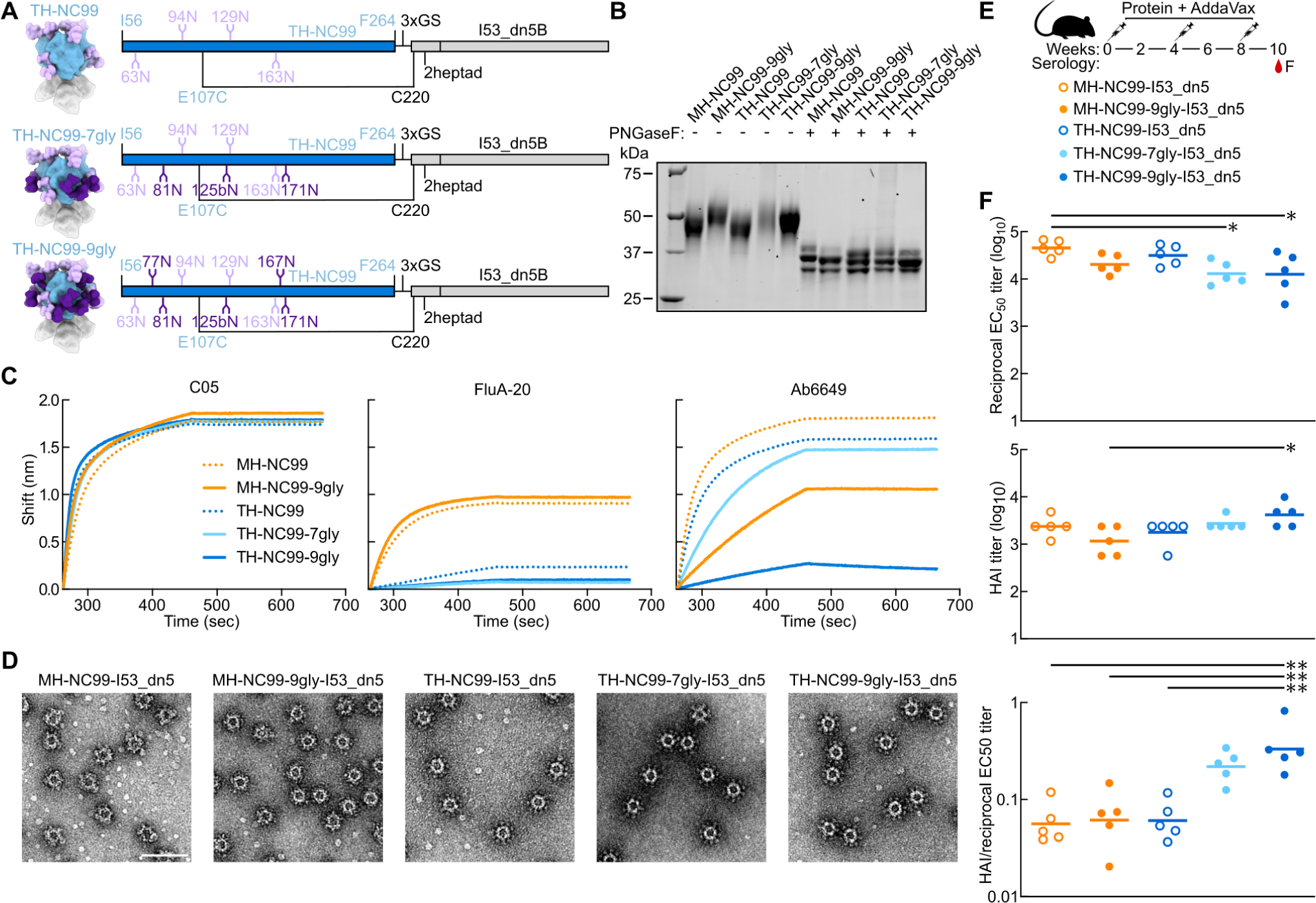
Design and Immunogenicity of Hyperglycosylated NC99 Trihead Nanoparticle Immunogens. (A) Model structures and gene diagrams for wild-type and hyperglycosylated NC99 triheads with wild-type glycans in light purple and glycan knock-ins in dark purple. NC99 HA numbering is in blue and trihead model numbering is in black. (B) Reducing SDS-PAGE of wild-type and hyperglycosylated NC99 monoheads and triheads without and with PNGaseF digestion. (C) BLI of wild-type and hyperglycosylated NC99 monoheads and triheads against C05, FluA-20, and Ab6649. (D) nsEM micrographs of hyperglycosylated NC99 monohead and trihead I53_dn5 nanoparticles. Scale bar = 100 nm. (E) Schematic illustrating mouse study timeline, immunizations, and serology timepoint. (F) Week 10 NC99-foldon trimer ELISA titers plotted as the reciprocal EC_50_ titer, hemagglutination inhibition (HAI) titers, and the ratio of HAI/reciprocal EC_50_ titers of hyperglycosylated NC99 monohead and trihead nanoparticles in BALB/c mice. Statistical significance was determined using one-way ANOVA with Tukey’s multiple comparisons test; ∗p < 0.05; ∗∗p < 0.01.

To quantify glycan occupancy at each site, all trihead components were analyzed using peptide mass spectrometry (Struwe et al. 2017; Stavenhagen et al. 2013). Glycan occupancy in TH-NC99 was high across all sites with the exception of N163, which was only about half occupied (45.1%; **Table S2**). The wild-type sites in TH-NC99-7gly showed similar occupancy, including half occupancy at N163 (44.3%). The engineered glycosylation sites in TH-NC99-7gly had high occupancy at N125b, about half occupancy at N81 (46.2%), and low occupancy at N171 (20.1%). The additional site at N125b was not observed as an isolated glycopeptide and could only be indirectly quantified by the peptide spanning the additional glycan site at N129. For TH-NC99-9gly, the additional N77 glycosylation site was only observed in a glycopeptide that also comprised N81 and showed either single (63%) or double (37%) occupancy. The last two sites in TH-NC99-9gly were only observed on a long glycopeptide containing three sequons. Based on the occupancy levels measured, it is likely that N167 is highly occupied (>90%) while N171 is less than 15% occupied, as this would be consistent with the occupancy levels observed for these sites in TH-NC99-7gly.

The monohead and trihead components were then combined with purified I53_dn5A pentamer *in vitro* at a 1:1 molar ratio to form icosahedral I53_dn5 nanoparticles displaying either 60 monohead monomers or 20 trihead trimers. Purification by SEC (**Figure S1C**) yielded pure, monodisperse preparations of nanoparticles according to SDS-PAGE, negative stain electron microscopy (nsEM), and dynamic light scattering (DLS) (**Figures 1D, S1D, and S1E**). Efficient assembly was observed for all trihead nanoparticles, as little residual component was observed during SEC purification (**Figure S1C**; peak at ∼17 mL). By contrast, substantial residual component was observed during SEC of the monohead nanoparticle assemblies, indicating less efficient assembly.

We evaluated the hyperglycosylated trihead nanoparticle immunogens in an initial immunogenicity study in mice. BALB/c mice were immunized with 1.5 μg of nanoparticle immunogen formulated with AddaVax at weeks 0, 4, and 8 **(Figure 1E)**. Strain-matched NC99 HA-binding antibody titers from serum collected at week 10 showed reduced binding in all hyperglycosylated groups compared to their wild-type counterparts **(Figure 1F)**. Conversely, NC99 hemagglutination inhibition (HAI) titers were highest in the TH-NC99-9gly group and lowest in the MH-NC99-9gly group. Plotting the ratio of HAI/binding titers revealed a trend towards a stepwise increase with increasing glycosylation in the trihead groups, suggesting a higher proportion of on-target receptor-blocking antibodies. Only the MH-NC99-9gly sera competed with FluA-20 binding in competition ELISAs, yet these sera showed the least amount of competition with C05 **(Figure S1F-S1G)**. These results suggest that hyperglycosylation refocused vaccine-elicited antibodies onto receptor-blocking epitopes in the case of the trihead immunogens, and onto the trimer interface in the case of the monohead immunogens.

### Design of Hyperglycosylated Trihead Antigens from Additional H1 HAs

We and others have recently reported that mosaic nanoparticle immunogens, which co-display multiple antigenic variants on the same nanoparticle surface, can induce broadly protective responses against related viruses by eliciting antibodies that target conserved epitopes (Kanekiyo et al. 2019; Boyoglu-Barnum et al. 2021; Cohen, Gnanapragasam, et al. 2021; Walls et al. 2021; Sliepen et al. 2022; Cohen et al. 2022; Brinkkemper et al. 2022). To enable mosaic trihead display as a potential route to enhancing breadth amongst H1 strains, we adapted the trihead design strategy to three other divergent H1s with unique antigenic properties: A/South Carolina/1/1918 (TH-SC18), A/Puerto Rico/8/1934 (TH-PR34), and A/Michigan/45/2015 (TH-MI15). We again made corresponding monohead antigens for comparison. These antigens were all connected to the I53_dn5B trimer using one heptad repeat of the GCN4-based coiled-coil, as this rigid linker length was found to yield optimal cross-reactive antibody responses in mice (Ellis et al., n.d.). The same disulfide bond in TH-NC99 between the base of the trihead and the coiled-coil linker was used, as well as similar stabilizing mutations at the trimer interface, although the amino acids used at positions 203 and 205 differed among strains (**Figure 2A** and **Table S1**). Glycan knock-in mutations were included in final designs for TH-PR34, at position 63, and for MH-SC18 and TH-SC18, at position 125b, which dramatically enhanced secretion and stability. Additional resurfacing mutations P26S and V84E in TH-PR34, as well as A198E in both TH-SC18 and TH-MI15, were also key to enhancing secretion.

**Figure 2.**
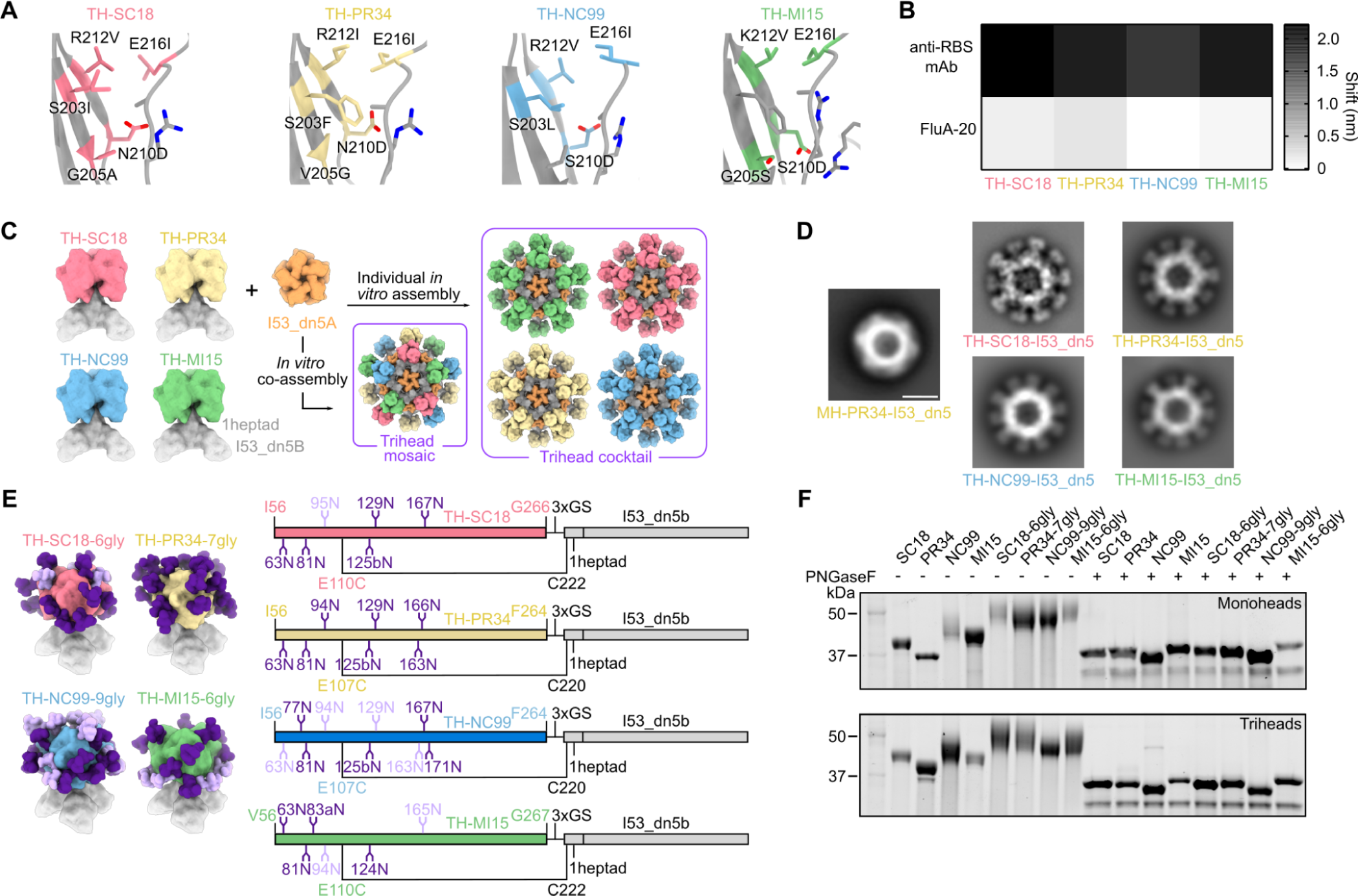
Design of Hyperglycosylated Trihead Antigens from Additional H1 Has. (A) Diagram of head trimer interfaces for TH-SC18, TH-PR34, TH-NC99, and TH-MI15, where mutated residues are colored and labeled. (B) BLI of trihead components against RBS-directed mAbs (5J8, anti-PR34, and C05) and FluA-20. (C) Schematic of TH-SC18, TH-PR34, TH-NC99, and TH-MI15 constructs and their *in vitro* assembly into mosaic or cocktail I53_dn5 nanoparticles. (D) nsEM 2D class averages of MH-PR34-I53_dn5 and trihead I53_dn5 nanoparticles. Scale bar = 25 nm. (E) Model structures and gene diagrams for hyperglycosylated triheads with wild-type glycans in light purple and glycan knock-ins in dark purple. Strain-specific H1 HA numbering is in respective HA strain color and trihead model numbering is in black. (F) Reducing SDS-PAGE of wild-type and hyperglycosylated monoheads and triheads without and with PNGaseF digestion.

All four triheads maintained binding to RBS-directed antibodies, with minimal FluA-20 binding by BLI, indicating trihead closure (**Figures 2B and S2A**). By contrast, monohead versions of each strain all showed high binding to both RBS antibodies and FluA-20. The trihead and monohead components were purified using SEC (**Figures S2B and S2C**) prior to *in vitro* assembly into I53_dn5 nanoparticles. We prepared a cocktail of nanoparticles by mixing together the four individually assembled trihead nanoparticles, as well as mosaic nanoparticles in which the four trihead components were mixed together prior to addition of I53_dn5A pentamer (**Figure 2C**). All nanoparticles were then purified using SEC and their purity and monodispersity were verified by SDS-PAGE, DLS, and nsEM (**Figures S3A-S3D**). All four monovalent trihead nanoparticles exhibited individually resolved trihead densities on the nanoparticle exteriors in nsEM averages, while the MH-PR34-I53_dn5 nanoparticle lacked visible antigen density due to the flexibility of the monoheads (**Figure 2D**). Taken together, the BLI, SEC, and nsEM averages indicate the formation of closed, relatively rigid trihead nanoparticle immunogens for all four strains.

As with TH-NC99-9gly, engineered glycans were used to mask epitopes outside the RBS in each H1 trihead. Individual glycans added to TH-SC18 were tested for their effects on trihead secretion before combination into one hyperglycosylated construct. These data were then used to guide hyperglycosylation of TH-PR34 and TH-MI15. TH-SC18-6gly has one wild-type and five engineered glycans, TH-PR34-7gly has zero wild-type and seven engineered glycans, and TH-MI15-6gly has two wild-type and five engineered glycans (**Figure 2E** and **Table S1**). All hyperglycosylated trihead components showed apparent increases in molecular weight by SDS-PAGE compared to their wild-type counterparts, while treatment with PNGase F resulted in nearly uniform migration of all constructs at a lower apparent molecular weight (**Figures 2F and S2C**). All hyperglycosylated trihead components maintained the desired antigenic profile of high anti-RBS antibody binding with minimal FluA-20 binding and had circular dichroism (CD) spectra that closely matched those of the corresponding non-hyperglycosylated triheads, indicating they retained their native structure (**Figures S4A-S4C**). The four hyperglycosylated trihead components were combined in an equimolar mixture and then co-assembled *in vitro* with the I53_dn5A pentamer to form a hyperglycosylated trihead mosaic nanoparticle that was purified by SEC (**Figure S3A**). SDS-PAGE, DLS, and nsEM of the purified assembly revealed monodisperse nanoparticles of the expected size and morphology (**Figures S3B-S3D**).

We compared the stabilities of the hyperglycosylated monohead and trihead components using hydrogen-deuterium exchange mass spectrometry (HDX-MS) and nano differential scanning fluorimetry (nanoDSF). Deuterium uptake profiles across MH-NC99-9gly and all four hyperglycosylated triheads were consistent with the antigens adopting similar conformations, although with some local differences (**Figure S5A**). For example, HDX-MS analysis of RBS peptides showed that TH-SC18-6gly has greater dynamics at the 190-helix and 220-loop relative to the other three strains of hyperglycosylated triheads (**Figures S5B-S5C**). Similarly, trimer interface peptides lying in the 220-loop and 200-loop in the TH-SC18-6gly were also less ordered, especially compared to TH-NC99-9gly and TH-PR34-7gly, which were the most ordered in these regions. Comparing MH-NC99-9gly and TH-NC99-9gly showed that the monohead antigen displayed substantially higher exchange than the trihead at the trimer interface in the 220-loop, as well as slightly elevated dynamics in other regions. Thermal denaturation monitored by SYPRO Orange fluorescence showed similar trends in stability, where TH-SC18-6gly had the lowest melting temperature (T_m_) and TH-NC99-9gly had the highest T_m_ among the triheads (**Figure S5D**). Additionally, three out of four hyperglycosylated monoheads had T_m_s ≥4°C lower than their trihead counterparts. Together, these analyses indicate that the trihead antigens are locally and globally more stable than monomeric RBDs in addition to being rigidly linked to the I53_dn5 nanoparticle scaffold.

### Design and Characterization of Hypervariable Trihead Immunogens

As a further test of the hypothesis that mosaic nanoparticle display focuses antibody responses on conserved epitopes, we designed a hypervariable antigen library featuring mutations within the RBS periphery. This is a particularly variable region in hemagglutinins from different influenza virus strains, and mutations in this region are a central driver of antigenic drift (**Figure 3A**) (Koel et al. 2013). We reasoned that co-display of a library of trihead variants with mutations in the RBS periphery may elicit fewer strain-specific antibodies in favor of responses targeting the conserved RBS. We constructed our library to recapitulate this variability by introducing naturally occurring mutations or those identified by deep mutational scanning into key hypervariable positions (**Figure 3A**) (Doud and Bloom 2016; Wu et al. 2017). We made four variants of each hyperglycosylated trihead, each comprising a unique combination of 2-10 amino acid mutations in the RBS periphery (**Figure 3B** and **Table S3**). Binding studies using BLI showed that the variants had distinct antigenic profiles as intended, with some mutations leading to a complete loss of binding to particular anti-RBS mAbs (**Figures 3C and S4A**). However, all RBS variants maintained minimal FluA-20 binding, exhibited the expected SEC elution profiles and apparent molecular weights on SDS-PAGE, and had similar CD spectra showing a mixture of alpha helix and beta sheet, indicating that they formed well-folded, closed triheads (**Figures 3C, S4C, and S6A-S6B**). The four base hyperglycosylated triheads and all of their variants were then pooled and co-assembled with I53_dn5A pentamer to generate a hyperglycosylated, hypervariable nanoparticle containing 20 unique trihead antigens (**Figures 3D**). This hypervariable trihead nanoparticle was purified by SEC and was monodisperse by SDS-PAGE, DLS, and nsEM (**Figures S3A-S3D**). Co-display of all trihead mosaic nanoparticles (TH-mosaic-I53_dn5, TH-hyperglycosylated-mosaic-I53_dn5, and TH-hypervariable-hyperglycosylated-mosaic-I53_dn5) was confirmed using sandwich BLI by comparison to the TH-cocktail-I53_dn5 immunogen (**Figure S6C**). The NC99-reactive mAb C05 was first loaded onto AR2G biosensors, followed by nanoparticle loading and sequential binding to a PR34-specific mAb (Sino Biological) and 5J8, which binds both MI15 and SC18. Although all nanoparticles bound to the immobilized C05 mAb, only the three mosaic nanoparticles showed subsequent binding to the PR34-specific mAb and 5J8. These data indicate that the mosaic nanoparticles co-display trihead antigens that bind all three antibodies, and that there is no detectable subunit exchange in the cocktail nanoparticle preparation. We note that the lower amount of hypervariable trihead nanoparticle loading and subsequent antibody binding is consistent with the individual hypervariable trihead component BLI, which demonstrated that some mutations within the RBS periphery abrogated specific mAb binding, particularly for the anti-PR34 mAb (**Figure 3C** and **S4A**).

**Figure 3.**
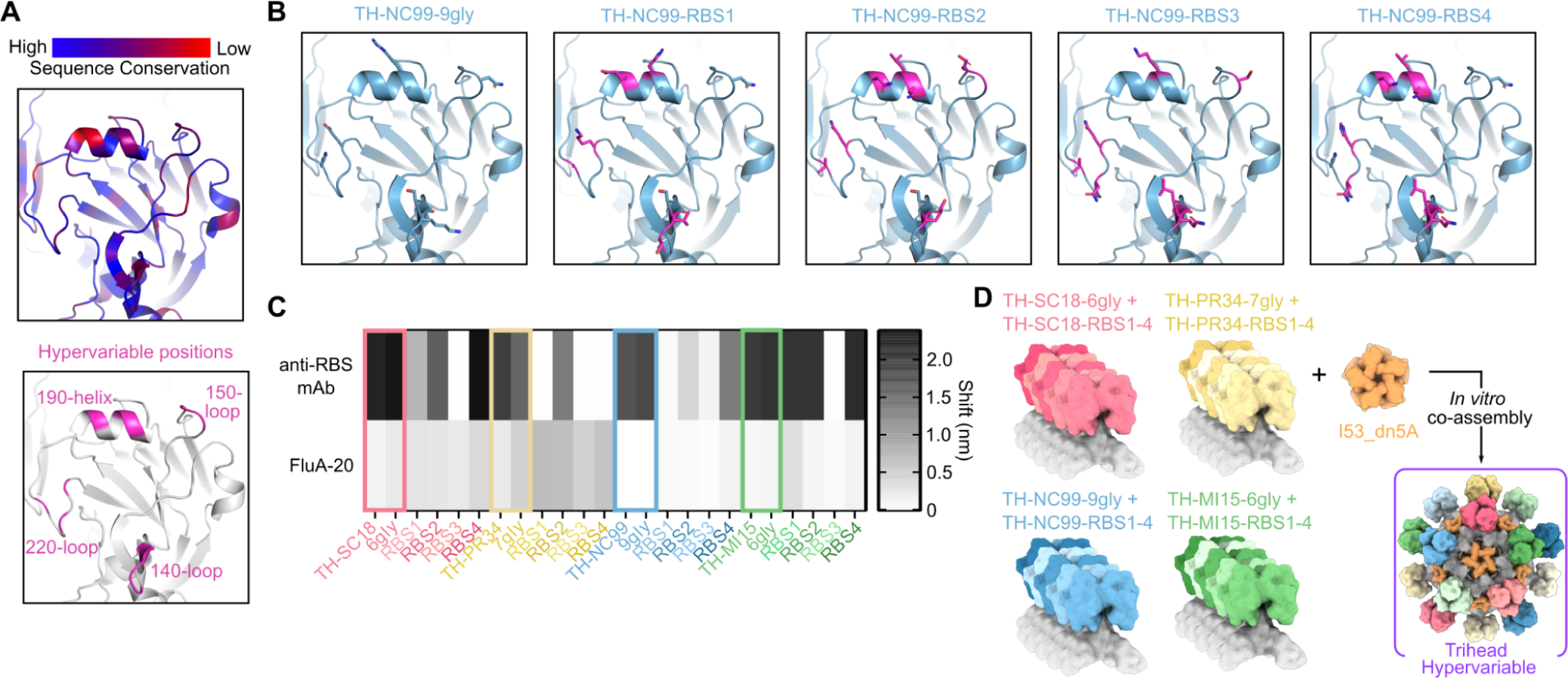
Design and Characterization of Hypervariable Trihead Immunogens. (A) Sequence conservation amongst 643 unique H1 sequences (top) and positions mutated in hypervariable library as dark pink (bottom) modeled on the NC99 HA structure (PDB: 7SCN). (B) TH-NC99-9gly wild-type and hypervariable variants modeled onto the NC99 HA structure (PDB: 7SCN), with all positions mutated in the library shown as sticks, wild-type residues in blue, and mutated residues in magenta. (C) BLI of triheads and hyperglycosylated triheads, with colored squares around these constructs, and trihead RBS variant components against RBS-directed mAbs (5J8, anti-PR34, and C05) and FluA-20. (D) Schematic of hypervariable trihead components and assembly into an I53_dn5 nanoparticle.

### Vaccine-elicited Antibody Responses in Rabbits Immunized with Monohead and Trihead Nanoparticles

We then evaluated our series of monohead and trihead nanoparticle immunogens in an immunogenicity study in New Zealand white rabbits. We chose rabbits because their long CDRH3 repertoire may facilitate the elicitation of antibodies that can penetrate into the RBS (Whittle et al. 2011; Ekiert et al. 2012; Sanders et al. 2015), and also because they permit the collection of sufficient serum to conduct a number of distinct serological analyses. Rabbits were immunized at weeks 0, 4, and 20 with 25 μg immunogen formulated with AddaVax (**Figure 4A**). Negligible binding titers at week 0 against a vaccine-matched MI15 HA-foldon trimer showed that there were no pre-existing anti-HA antibodies in these animals (**Figure S7A-B**). Evaluation of vaccine-matched (NC99 and MI15) serum antibody binding, HAI, and microneutralization using sera obtained at weeks 6 and 22 revealed several differences between the groups. First, the cocktail and mosaic monohead groups consistently had lower binding, HAI, and neutralization titers than all other groups at week 6, although the differences were not always statistically significant (**Figures 4B-D**). Interestingly, the hyperglycosylated monohead mosaic elicited antibody responses that were significantly higher than the other monohead groups and comparable to the trihead immunogens. At week 22, the monohead cocktail group had significantly lower NC99-neutralizing activity than all other groups, as well as NC99 HAI titers that trended lower (**Figures 4E and S5C-D**). By contrast, the monohead and trihead mosaic groups had significantly lower MI15-neutralizing activity than most of the other groups (**Figure 4E**). Taken together, the data show that the monohead cocktail and mosaic nanoparticle immunogens were generally less immunogenic than the other groups, while the hyperglycosylated and hypervariable nanoparticles consistently elicited potent vaccine-matched responses.

**Figure 4.**
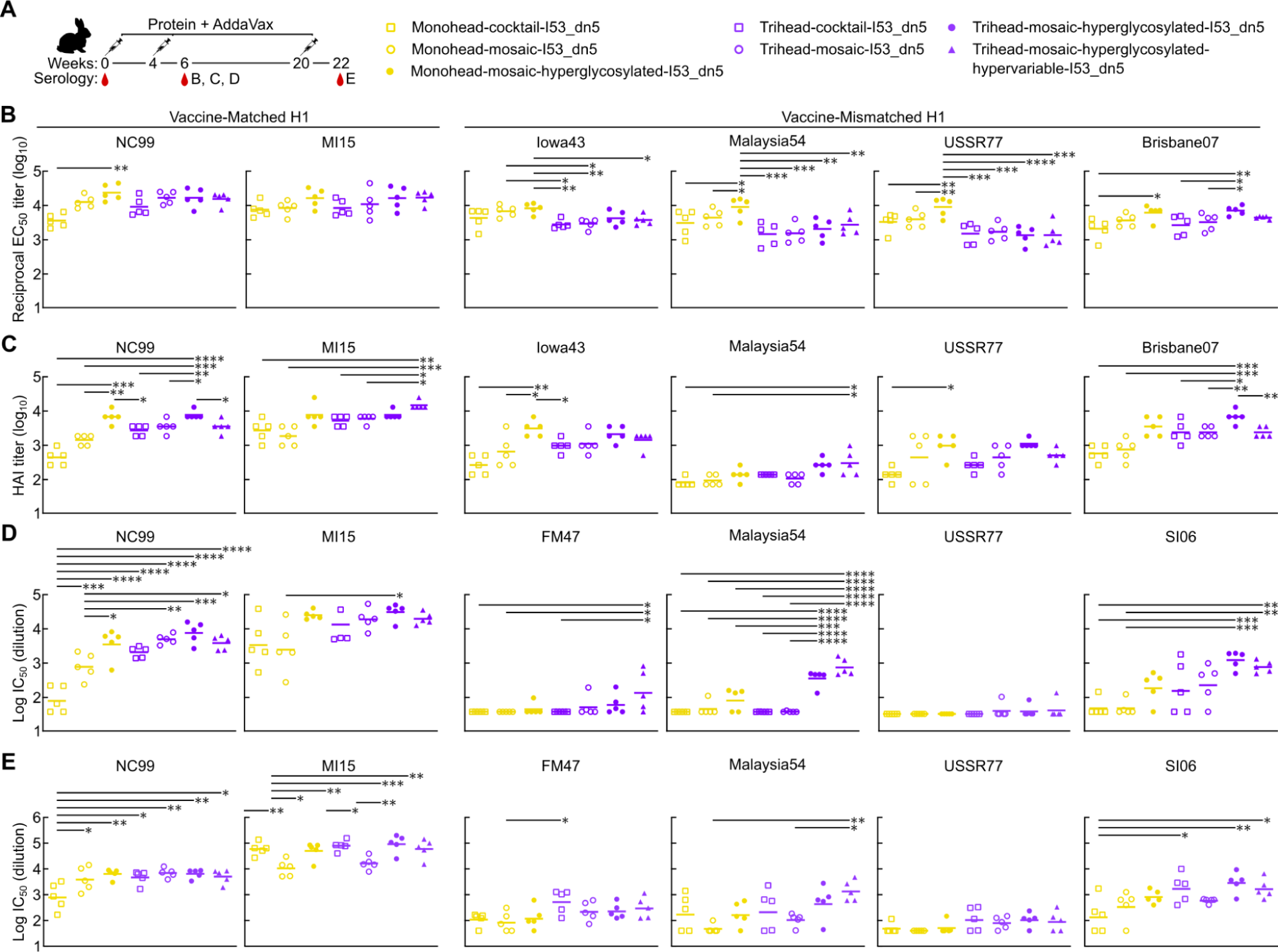
Vaccine-elicited Antibody Responses in Rabbits Immunized with Monohead and Trihead Nanoparticles. (A) Hypervariable trihead nanoparticle rabbit immunization schedule and groups. (B-D) B. ELISA binding titers, C. HAI titers, and D. Microneutralization titers in immune sera at week 6. (E) Microneutralization titers at week 22. Statistical significance was determined using one-way ANOVA with Tukey’s multiple comparisons test; ∗p < 0.05; ∗∗p < 0.01; ∗∗∗p < 0.001; ∗∗∗∗p < 0.0001.

Across four vaccine-mismatched H1 strains (see **Table S1**), the hyperglycosylated monohead group had the highest binding titers of all groups, both at weeks 6 and 22 (**Figures 4B and S7C**). The hyperglycosylated monohead also induced relatively high levels of HAI, however it elicited lower neutralizing activity than the hyperglycosylated trihead groups for all mismatched H1 strains at both weeks 6 and 22, although this was not always significant (**Figures 4C-E and S7D**). The relatively low level of neutralizing activity despite high binding titers indicates that the hyperglycosylated monohead nanoparticle elicited a high proportion of non-neutralizing antibodies, likely targeting the trimer interface. Interestingly, glycan masking of antigens has been shown to alter where on the antigen the majority of responses are being directed, but does not change the overall magnitude of the response (Duan et al. 2018; Bajic et al. 2019). By contrast in this study, the hyperglycosylated monohead mosaic outperformed its non-hyperglycosylated comparator group in every comparison, including overall binding titers across the strains tested, although not always with statistical significance. Alongside the hyperglycosylated monohead, the hyperglycosylated trihead group had the highest mismatched HAI responses against most strains tested (Iowa43, USSR77, and Brisbane07, see **Table S2**) with the exception of Malaysia54, where the hypervariable trihead group induced the highest HAI titers. This trend in HAI was mirrored in the microneutralization assays, where the highest neutralizing titers were obtained from the two hyperglycosylated trihead groups at weeks 6 and 22 for Malaysia54, and at week 6 for SI06. For the mismatched viruses FM47 and USSR77, neutralizing responses were low across all groups at week 6, with the exception of measurable activity in the hypervariable trihead group. However, by week 22 all trihead groups neutralized these viruses more potently than all monohead groups. Taken together, the consistently high vaccine-mismatched HAI and neutralization obtained from the two hyperglycosylated trihead groups suggests that stable trihead closure and hyperglycosylation both contributed towards eliciting superior immune responses.

### Epitope Mapping of Vaccine-Elicited Antibody Responses

We next sought to determine the epitope specificities of the serum antibodies elicited by each vaccine (**Figure 5A**). We first compared RBS knockout probes NC99-L194W and NC99-T155N/K157T to wild-type NC99 as ELISA antigens to assess the fraction of the antibody response in each group directed at the RBS.

**Figure 5.**
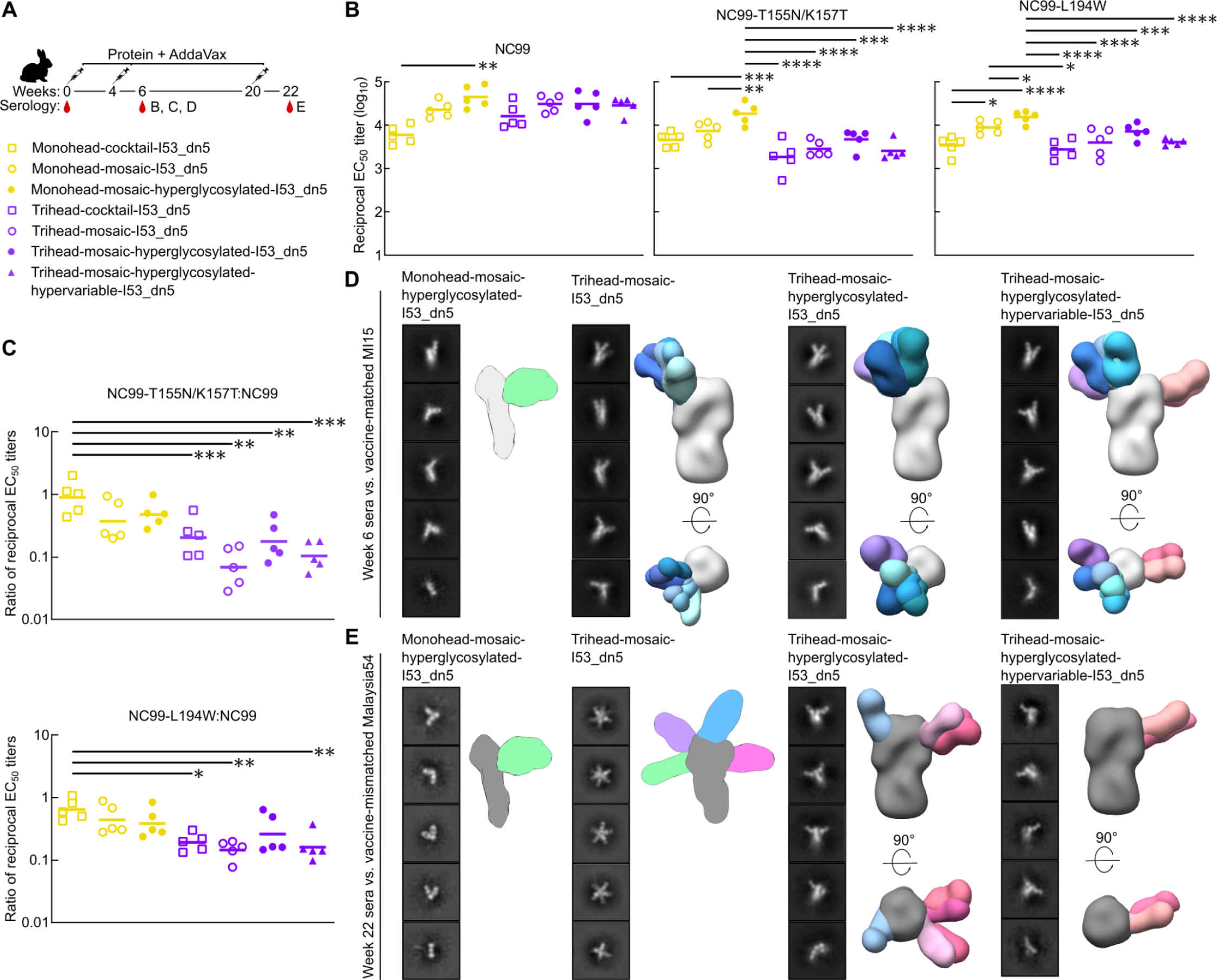
Epitope Mapping of Vaccine-elicited Antibody Responses. (A) Hypervariable trihead nanoparticle rabbit immunization schedule and groups. (B) ELISAs using NC99 probes against week 6 rabbit study serum. NC99 ELISA is the same as in Figure 4B. (C) Ratio of NC99 probes to NC99 binding titers in panel B. (D) Representative 2D class averages of week 6 serum from four groups in rabbit study against strain-matched MI15. Hyperglycosylated monohead group has a cartoon schematic of a likely 3D model, while all other groups are composite 3D models of ns-EMPEM analysis. (E) Representative 2D class averages of week 22 serum from four groups in rabbit study against strain-mismatched Malaysia54. Hyperglycosylated monohead and mosaic trihead groups have a cartoon schematic of their likely 3D models, while other groups are composite 3D models of ns-EMPEM analysis. Statistical significance was determined using one-way ANOVA with Tukey’s multiple comparisons test; ∗p < 0.05; ∗∗p < 0.01; ∗∗∗p < 0.001; ∗∗∗∗p < 0.0001.

NC99-L194W mutates a key highly conserved residue within the RBS pocket, introducing steric bulk that knocks out only antibodies that bind directly to the sialic acid binding pocket, while NC99-T155N/K157T introduces an N-linked glycan that more broadly prevents binding of RBS-directed antibodies (Ellis et al., n.d.). Serum antibody binding to both RBS knockout probes was considerably lower than to wild-type NC99 in all trihead groups (binding ratios of 0.08-0.64 for NC99-L194W and 0.03-0.65 for NC99-T155N/K157T), while the differences in relative binding in the monohead groups was smaller (binding ratios of 0.23-1.07 and 0.19-1.07, respectively) (**Figure 5B-C**). We attribute the greater reduction in binding using the T155N/K157T probe compared to the L194W to the fact that the glycan knock-in probe will interfere with antibodies targeting a larger area of the antigen. These results clearly indicate that the trihead immunogens elicited substantially more RBS-directed responses than the monohead immunogens.

We also used nsEM polyclonal epitope mapping (ns-EMPEM; (Bianchi et al. 2018; Han et al. 2021)) to directly visualize where vaccine-elicited serum antibodies bound HA. MI15 strain-matched ns-EMPEM using week 6 serum only detected a small fraction of the antibodies elicited by the hyperglycosylated monohead mosaic bound to the RBS of intact HA trimers, with most Fabs observed to be bound to HA monomers (**Figure 5D**). A substantial amount of these Fabs likely bind the trimer interface, which has been shown to disrupt trimerization, and indeed our class averages closely resemble those originally reported for FluA-20 (Bangaru et al. 2019). By contrast, all three mosaic trihead immunogens elicited antibodies that bound mostly to the RBS of intact HA trimers, though a small fraction of Fab bound to monomeric HA was also observed in these groups. We were able to discern RBS-targeting antibody classes with several slightly different angles of approach. Notably, the hypervariable trihead group also exhibited density for Fabs binding to the side of the MI15 HA head. This may be attributed to the lack of a glycan in TH-PR34-7gly, TH-MI15-6gly, and TH-SC18-6gly near residues 119-122 (SC18 numbering), in addition to the low occupancy of the N-linked glycan introduced at position 171 in TH-NC99, resulting in inefficient masking at this site. We also performed ns-EMPEM against the vaccine-mismatched Malaysia54 HA using week 22 sera to determine which epitopes were targeted by cross-reactive vaccine-elicited antibodies. We again observed predominantly Fab-bound HA monomers in the hyperglycosylated monohead group, indicating a high proportion of trimer interface-directed responses (**Figure 5E**). Week 22 trihead mosaic serum also revealed only Fab-bound HA monomers, but in contrast to the hyperglycosylated monohead group, the vast majority were bound to a multitude of Fabs targeting various epitopes in the head domain. Finally, in both the hyperglycosylated and hypervariable mosaic trihead sera we observed Fabs bound only to trimeric Malaysia54 HA, mostly targeting the side of the head, in contrast to the predominantly RBS-directed antibodies seen in these sera at week 6 against the vaccine-matched MI15.

We draw several conclusions from these RBS knockout probe binding and ns-EMPEM data. First, both sets of data suggest that monomeric RBD antigens elicit a high proportion of trimer interface-directed antibodies, while the closed, stabilized trihead antigens elicit antibodies predominantly targeting the RBS. Second, the week 22 mismatched ns-EMPEM suggests that hyperglycosylation further decreases the elicitation of trimer interface-directed antibodies by trihead immunogens. Third, the cross-reactive responses obtained with the monohead antigens largely derived from trimer interface-directed antibodies, explaining why vaccine-mismatched neutralization is low. Finally, the cross-reactive responses obtained with the hyperglycosylated trihead immunogens predominantly target a site on the side of the head near the previously identified lateral patch (Raymond et al. 2018) rather than the RBS, especially for the hypervariable immunogen that contains numerous mutations in the RBS periphery.

## DISCUSSION

Here we show that the initial NC99 trihead design strategy described in the accompanying manuscript (Ellis et al., n.d.) can be applied to several divergent H1 HAs, and that these trihead mutations improve the potency and breadth of vaccine-elicited antibody responses. The ability of design approaches or specific mutations to generalize across viral families is an important criterion in antigen design that has led to the generation of a number of promising antigen platforms. For example, “stabilized-stem” antigens based on influenza HA were made for both group 1 and 2 HAs, and when displayed on ferritin nanoparticle immunogens they elicited bnAb responses in animals as well as humans (Yassine et al. 2015; Corbett et al. 2019; Moin et al. 2022; Andrews et al. 2023; Darricarrère et al. 2021; Widge et al. 2023). We recently reported the stabilization of the closed tetrameric state of several different influenza neuraminidases, using an approach that was largely inspired by homology-guided mutations (Ellis et al. 2022). For several class I fusion glycoproteins, strategic use of proline mutations to stabilize the prefusion conformation (or destabilize the postfusion conformation) has proven to be a widely applicable design strategy (Sanders and Moore 2021). Proline mutations have been key in the generation of antigens that elicit potent neutralizing antibody responses against several viruses including HIV (Sanders et al. 2013, 2015) and RSV (Krarup et al. 2015), and provided an antigen platform that enabled rapid pandemic response vaccines against SARS-CoV-2 (Pallesen et al. 2017; Hsieh et al. 2020; Wrapp et al. 2020). The generalizability and improved immunogenicity we report here and in the accompanying manuscript (Ellis et al., n.d.) establish triheads as a promising new antigen platform for influenza vaccine design.

The higher HAI and neutralizing antibody responses we obtained with hyperglycosylated antigens are consistent with previous studies that have established hyperglycosylation as an effective antigen design strategy, while also raising mechanistic questions that motivate further studies. Our detailed antigenic characterization clearly revealed that several epitopes were successfully masked by the additional glycans in our hyperglycosylated immunogens, reducing the overall peptidic surface that can be targeted by antibodies. Nevertheless, in our rabbit study the hyperglycosylated monohead immunogen elicited higher binding titers across various H1 strains than either non-hyperglycosylated monohead immunogen (i.e., cocktail or mosaic). Though we did not assess glycan composition, a possible explanation for this discrepancy is that the potential presence of high-mannose glycans in the hyperglycosylated immunogens may drive better antigen trafficking to lymph nodes and B cell follicles, as demonstrated in recent studies from the Irvine lab (Read et al. 2022; Tokatlian et al. 2019). This effect could result in overall increases in the magnitude or quality of the vaccine-elicited antibody response that are independent of any potential redirection of vaccine-elicited antibody responses to target epitopes due to glycan masking. It is likewise possible that the hyperglycosylated immunogens are less susceptible to proteolytic degradation *in vivo*, which could also increase immunogenicity overall (Cirelli et al. 2019; Aung et al. 2023). These potential mechanisms are also consistent with the antibody responses elicited by our hyperglycosylated and non-hyperglycosylated trihead immunogens, where the former elicited higher HAI and neutralizing activity despite effectively no changes in binding antibody titers. It therefore appears that hyperglycosylation “focused” the antibody response on target epitopes near the receptor binding site. Several previous studies have shown that hyperglycosylation tends to increase the proportion—but not the overall magnitude—of on-target antibodies (Duan et al. 2018; Bajic et al. 2019). In fact, overall vaccine-elicited antibody titers can be reduced by epitope masking (Weidenbacher and Kim 2019). One potential explanation for our observation of clear increases in HAI and neutralizing activity may be an increase in overall immunogenicity accompanied by a reduction in off-target antibody responses. Glycan-dependent effects on trafficking and stability may account for the former, while the epitope masking provided by the glycans could account for the latter. Our EMPEM data in aggregate support suppression of non-RBS responses by hyperglycosylation, with the caveat related to under-occupancy at position 171 of the hyperglycosylated NC99 trihead as noted above. In summary, we observed a focusing-like effect from our hyperglycosylated trihead immunogens, but which of several potential mechanisms was primarily responsible remains to be determined. Future studies on the contributions of various mechanisms to the overall response will benefit from the use of systematic series of immunogens like those described here, in the accompanying manuscript (Ellis et al., n.d.), and in previous work (Read et al. 2022; Kato et al. 2020; Abbott et al. 2018).

In contrast to trihead closure and hyperglycosylation, the effects of mosaic nanoparticle display and hypervariable antigen design were less clear in our experiments. In the first study of mosaic nanoparticle immunogens, a significant advantage in eliciting cross-reactive B cell responses was observed when presenting HA RBDs on ferritin nanoparticles in a mosaic array compared to either a cocktail or sequential immunization regimen (Kanekiyo et al. 2019). In this work, we observed more strain-specificity in vaccine-matched responses elicited by the monohead cocktail group compared to the monohead mosaic group. However, we did not observe any significant differences between other cocktail vs. mosaic comparisons for either monohead and trihead immunogens. The addition of the hypervariable RBS periphery also did not show any significant differences compared to the hyperglycosylated trihead group. There are several differences between this study and that of Kanekiyo and colleagues that could account for why mosaics were only superior in the latter, including animal model, HA strain compositions, nanoparticle size and valency, T cell epitope content, monomeric RBDs vs. trihead antigens, and flexible vs. rigid attachment to the nanoparticle scaffold. One noticeable similarity between the two studies is that the cross-neutralizing responses we observed seem to derive from antibodies directed against the side of the HA head that bind an epitope similar to that of 441D6 (Kanekiyo et al. 2019). Although it is likely that glycan under-occupancy at this site is partially responsible, it is intriguing to speculate that BCR cross-linking by mosaic nanoparticles could explain the boost in relatively rare cross-reactive antibodies against this epitope in both studies. Additional studies that more rigorously characterize this epitope on the side of the head and the antibody responses elicited against it could inform future vaccine design efforts.

In conclusion, we have shown that trihead nanoparticle immunogens are a promising platform for next-generation influenza vaccine design. Furthermore, our evaluation of multiple layers of immune focusing techniques provides a roadmap for efforts using this and other antigen platforms to develop safe and effective vaccines against a variety of pathogens.

## METHODS

### Gene Expression and Protein Purification

All HA constructs used in this study included the Y98F mutation (Whittle et al. 2014) and were codon-optimized for human cell expression and made in the CMV/R vector (Barouch et al. 2005) by Genscript with a C-terminal hexahistidine affinity tag. PEI MAX was used for transient transfection of HEK293F cells. After four days, mammalian cell supernatants were clarified via centrifugation and filtration. Monohead and trihead components and HA foldons were all purified using IMAC. 1 mL of Ni^2+^-sepharose Excel or Talon resin was added per 100 mL clarified supernatant along with 5 mL of 1 M Tris, pH 8.0 and 7 mL of 5 M NaCl and left to batch bind while shaking at room temperature for 30 min. Resin was then collected in a gravity column, washed with 5 column volumes of 50 mM Tris, pH 8.0, 500 mM NaCl, 20 mM imidazole, and protein was eluted using 50 mM Tris, pH 8.0, 500 mM NaCl, 300 mM imidazole. Further component purification was done using SEC on a Superdex 200 Increase 10/300 gel filtration column equilibrated in 25 mM Tris, pH 8.0, 150 mM NaCl, 5% glycerol. HA-ferritin nanoparticles used in HAI assays were purified as described previously (Kanekiyo et al. 2013).

Expression and purification of the I53_dn5A pentamer component from *E. coli* was carried out as previously described (Boyoglu-Barnum et al. 2021). Assembly of trihead-I53_dn5 nanoparticles was carried out by mixing purified HA trihead-I53_dn5B and pentameric I53_dn5A components together *in vitro* at a 1:1 molar ratio at 15-40 μM final concentrations. Nanoparticles were left to assemble for 30 min at room temperature with rocking. Nanoparticles were then purified using SEC on a Superose 6 Increase 10/300 gel filtration column equilibrated in 25 mM Tris, pH 8.0, 150 mM NaCl, 5% glycerol.

Following purification, nanoparticle quality and concentration was first measured by UV-vis spectroscopy. Nanoparticle polydispersity and purity was then assessed using SDS-PAGE, DLS, and nsEM. Finally, endotoxin levels were measured using the LAL assay, with all immunogens used in animal studies containing less than 100 EU/mg in the final dose. Final immunogens were flash-frozen using liquid nitrogen and stored at −80°C.

### Bio-layer Interferometry (BLI)

BLI was carried out using an Octet Red 96 system, at 25°C with 1000 rpm shaking. Anti-HA antibodies were diluted in kinetics buffer (PBS with 0.5% serum bovine albumin and 0.01% Tween) to a final concentration of 10 μg/mL before loading onto protein A biosensors (Sartorius) for 200 s. Monohead and trihead components were diluted to 500 nM in kinetics buffer and their association was measured for 200 s, followed by dissociation for 200 s in kinetics buffer alone.

### Sandwich BLI

AR2G biosensors (Sartorius) were first activated by the addition of freshly mixed 20 mM 1-Ethyl-3-(3-dimethylaminopropyl)carbodiimide and 10 mM Sulfo-N-hydroxysulfosuccinimide. C05 mAb at 5 μg/mL in 10 mM acetate buffer, pH 5.0 was then loaded, followed by quenching with 1 M ethanolamine, pH 8.5. Nanoparticles at 30 μg/mL in kinetics buffer (PBS with 0.5% serum bovine albumin and 0.01% Tween) were then loaded, followed by a baseline step in kinetics buffer before subsequent association and dissociation of anti-PR34 and then 5J8, both at 50 nM in kinetics buffer.

### Negative Stain Electron Microscopy

3.5 µl of 70 µg/mL nanoparticles were applied to glow-discharged 400-mesh carbon-coated grids (Electron Microscopy Sciences) and stained with 2% (wt/vol) uranyl formate. Data were collected using EPU 2.0 on a 120 kV Talos L120C transmission electron microscope (Thermo Scientific) with a BM-Ceta camera. CryoSPARC (Punjani et al. 2017) was used for CTF correction, particle picking and extraction, and 2D classification.

### Dynamic Light Scattering

DLS was carried out on an UNcle (UNchained Labs) at 25°C. 10 acquisitions of 5 sec each were acquired for each spectrum. Protein concentration (ranging from 0.1-1 mg/mL) and buffer conditions were accounted for in the software.

### Glycan Occupancy Quantitation

Glycan occupancy quantitation was performed using peptide mass spectrometry to determine the relative abundance of peptides in the non-glycosylated and de-glycosylated states after full deglycosylation using N-glycanase (Stavenhagen et al. 2013). Each construct was combined with guanidine hydrochloride and DTT (6 M and 20 mM final concentration, respectively) and boiled for 30 minutes. Cysteines were then alkylated with the addition of 40 mM iodoacetamide and incubated in the dark for 1 hour and quenched with another addition of 20 mM DTT. Samples were diluted 6-fold in 10 mM Tris pH 8.0, 1 mM CaCl_2_ and treated with N-glycanase (New England Biolabs) for 1 hour at 37°C.

The samples were then split into two and treated with either LysC or GluC proteases (ThermoScientific, 1:20 protease:substrate molar ratio) overnight at 37°C. Digestions were quenched with the addition of 0.25% formic acid. Peptides were trapped and desalted using C18 spin columns (ThermoScientific) using the manufacturer’s suggested protocol and dried by speedvac. Purified peptides were resuspended in 20 µL of 0.1% formic acid. LC-MS analysis was performed on an Thermo EASYnLC coupled to a Thermo Orbitrap Fusion operating in data-dependent mode using EThcD fragmentation. Peptides were resolved over a pulled 30 µm ID silica tip packed with Reprosil-Pur 120 C18-AQ, 5 µm (ESI Source Solutions) using a linear gradient of 5-30%B (A: 0.1% formic acid; B: acetonitrile with 0.1% formic acid) over 90 minutes at a flowrate of 300 nL/min. LC-MS data were analyzed with Byonic (Protein Metrics Inc.) with a score cutoff of 100 and quantitative analysis was performed using Skyline (Pino et al. 2020).

### Circular Dichroism

CD measurements were carried out on a JASCO J1500 spectrometer at 25°C, using 1 mm path-length cuvette, at wavelengths from 200-260 nm. Proteins were measured at 0.2-0.3 mg/ml in TBS buffer.

### Hydrogen Deuterium Exchange Mass Spectrometry

40 pmol of each protein per timepoint were incubated in deuterated buffer (20 mM PBS, 85% D2O, pH∗ 7.48) for 3 s, 1 min, 30 min, and 22 hrs at room temperature (23°C). The reaction was stopped by diluting 1:1 in ice-cold quench buffer (200 mM tris(2-chlorethyl) phosphate (TCEP), 4 M urea, 0.2% formic acid) to a final pH of 2.495. Samples were immediately flash frozen in liquid nitrogen and stored at −80°C prior to analysis. As previously described, online nepenthesin-2 (Nepenthesin-2 protease column, 2.1 x 20 mm, purchased from POROS) digestion was performed and analyzed by LC-MS-IMS utilizing a Waters ACQUITY UPLC CSH C18 VanGuard column (130 Å, 1.7 μm, 1mm x 100 mm) and a Waters Synapt G2-Si Q-TOF mass spectrometer (Verkerke et al. 2016). A custom HDX cold box maintained the protease digestion at 4°C and the LC plumbing at 0°C throughout the 15 minute gradient (Watson et al. 2021). Waters MSe data was collected on the Waters Synapt G2-Si Q-TOF and was processed using Byonic (Version 3.8, Protein Metrics Inc.) to obtain a peptide reference list for each construct. Some homologous peptides were determined by aligning sequences using Clustal-Omega and calculating theoretical monoisotopic m/z values using ExPASy PeptideMass. Percent exchange values were calculated with theoretical total deuteration profiles produced by HD-Examiner (version 3.3, Seirra Analytics). An internal exchange standard (Pro-Pro-Pro-Ile [PPPI]) was included in each reaction to control for variations in ambient temperature during the labeling reactions. Data for each timepoint was collected in duplicate and error bars were plotted using one standard deviation. Back-exchange was determined by comparing experimental totally deuterated spectra to theoretical totally deuterated spectra. The average back exchange across all peptides was determined to be 31.6%. TH-NC99-9gly data was collected separately from the rest of the triheads and at a higher cone voltage (150 V) which reduced peptide signal at higher charge states (3+, 4+). To control for increases in back exchange from the higher cone voltage, a 3s and 1m control (TH-SC18-RBS2) was run at both voltages (40V and 150V) producing an average difference of 2.6% which is smaller than the differences we observe in the data.

### NanoDSF

All proteins were formulated at 0.5 mg/mL in 25 mM Tris, pH 8.0, 150 mM NaCl, 5% glycerol and then mixed at 9 volumes to 1 volume of 200× concentrate SYPRO orange (Thermo Fisher) diluted in the same buffer. NanoDSF to determine melting temperatures was carried out on an UNcle (UNchained Labs) by measuring the integration of fluorescence emission spectrum during a thermal ramp from 25°C to 95°C, with a 1°C increase in temperature per minute.

### HA Sequence Conservation Plot

643 unique H1 HA sequences were downloaded from the Influenza Research Database (https://legacy.fludb.org/). Sequence conservation amongst these was then plotted on HA using PyMOL (The PyMOL Molecular Graphics System, Version 2.0, Schrödinger, LCC).

### Immunization

Female BALB/c mice (Jackson Laboratories) were immunized intramuscularly with 1.5 μg purified nanoparticle immunogen in 100 μl (50 μl in each hind leg) of 50% (v/v) mixture of AddaVax adjuvant (Invivogen, San Diego, CA) (**Figure 1 and S1)**. For sera collection, mice were bled via submental venous puncture 2 weeks following each inoculation. Serum was isolated from hematocrit via centrifugation at 2,000 g for 10 min, and stored at −80°C until use. Female New Zealand white rabbits weighing approximately 4 kg were immunized intramuscularly with 25 μg purified nanoparticle immunogen in 1 ml (500 μl in each quadricep) of 50% (v/v) mixture of AddaVax adjuvant (**Figure 4, 5 and S7**).

### ELISA

HA-foldon trimers were added to 96-well Nunc MaxiSorp plates (Thermo Scientific) at 5.0 μg/mL with 50 μL per well and incubated for 1 hour. Blocking buffer composed of Tris Buffered Saline Tween (TBST: 25 mM Tris pH 8.0, 150 mM NaCl, 0.05% (v/v) Tween20) with 5% Nonfat milk was then added at 200 μl per well and incubated for 1 hour. Next plates were washed, with all washing steps consisting of 3x washing with TBST using a robotic plate washer (Biotek). 5-fold serial dilutions of serum starting at 1:100 were made in blocking buffer, added to plates at 50 μl per well, and incubated for 1 hour. Plates were washed again before addition of 50 μl per well of either anti-mouse or anti-rabbit HRP-conjugated goat secondary antibody (CellSignaling Technology) diluted 1:2,000 in blocking buffer and incubated for 30 minutes. All incubations were carried out with shaking at room temperature. Plates were washed a final time, and then 100 μl per well of TMB (3,3′,5′,5-tetramethylbenzidine, SeraCare) was added for 2 minutes, followed by quenching with 100 μl per well of 1 N HCl. Reading at 450 nm absorbance was done on an Epoch plate reader (BioTek).

### Competition ELISA

Competition ELISAs were performed in the same manner as the above ELISA protocol with some modifications as follows. For competition with FluA-20, 5-fold serial serum dilutions were made starting at 1:10, and for competition with C05, 3-fold serial serum dilutions were made starting at 1:5. Serum was left to incubate for 30 minutes, followed by addition of 50 μl of 0.1 μg/ml competitor antibody in blocking buffer and incubation for 45 minutes. After washing, anti-human HRP-conjugated goat secondary antibody (Southern Biotech) was added at 20,000x at 50 μl per well and incubated for 30 minutes.

### HAI

Serum was inactivated using receptor-destroying enzyme (RDE) II (Seiken) in PBS at a 3:1 ratio of RDE to serum for 16 hours at 37°C, followed by 40 minutes at 56°C. Inactivated serum was serially diluted 2-fold in PBS in V-bottom plates at 25 μl per well. 25 μl HA-ferritin nanoparticles at 4 hemagglutinating units were then added to all wells and incubated at room temperature for 30 min (Whittle et al. 2014). Lastly, 50 μl of 10-fold diluted turkey red blood cells (Lampire) in PBS was added to each well. Hemagglutination was left to proceed for at least 1 hour before recording HAI titer.

### Reporter-based microneutralization assay

Influenza A reporter viruses were made as previously described (Creanga et al. 2021). Briefly, H1N1 viruses were made with a modified PB1 segment expressing the TdKatushka reporter gene (R3ΔPB1), rescued, and propagated in MDCK-SIAT-PB1 cells in the presence of TPCK-treated trypsin (1 μg ml−1, Sigma) at 37 °C. Virus stocks were stored at −80 °C and were titrated before use in the assay. Rabbit sera was treated with receptor destroying enzyme (RDE II; Denka Seiken) and heat-inactivated before use in neutralization assays. 384 well plates (Greiner) were pre-seeded with 1.0 × 10^5 MDCK-SIAT1-PB1 cells and incubated overnight. Immune sera or monoclonal antibody controls (CR8071 and CR9114 (Dreyfus et al. 2012)) were serially diluted and incubated for 1 h at 37 °C with pre-titrated virus (A/Fort Monmouth/1/1947, A/Malaysia/302/1954, A/New Caledonia/20/1999, A/USSR/90/1977, A/Solomon Islands/3/2006, A/Michigan/45/2015). Serum-virus mixtures were then transferred in quadruplicate onto the pre-seeded 384 well plates and incubated at 37 °C for 18-26 hours. The number of fluorescent cells in each well was counted automatically using a Celigo image cytometer (Nexcelom Biosciences). IC50 values, defined as the serum dilution or antibody concentration that gives 50% reduction in virus-infected cells, were calculated from neutralization curves using a four-parameter nonlinear regression model.

### EMPEM

1 mL rabbit serum was diluted 3x in PBS and incubated overnight with 1 ml packed rProtein A Sepharose Fast Flow resin (Cytiva). The FT was removed using gravity purification and resin was washed with 20 CVs PBS. IgGs were eluted by incubating resin for 20 min with 1 CV of 0.1 M glycine, pH 2.5, repeated twice. Elutions were neutralized with 1 M Tris, pH 8.0 to a final concentration of 50 mM. IgGs were then buffer exchanged into PBS and concentrated to 250 μl for digestion into fabs. Fab digestion was carried out by adding in 250 μl freshly made 2x digestion buffer (40 mM NaPO4 pH 6.5, 20 mM EDTA, 40 mM Cysteine) and 500 μl papain resin in 500 μl 1x digestion buffer, and incubated with shaking at 37°C for 16 hours. Papain digestion reaction was centrifuged and supernatant containing fabs was collected and filtered. Papain was subsequently washed with 1 CV 20 mM Tris, pH 8.0 and this supernatant was added to the first. Digested sera was then purified by SEC on a Superdex 200 Increase 10/300 GL column. Purified fabs were then concentrated to 50 μl, mixed with 50-fold molar excess of HA-foldon trimers, and incubated for 16-20 hours at room temperature with gentle rocking. Immune complexes were purified by SEC on a Superdex 200 Increase 10/300 GL column and used in EM.

nsEM data were collected using EPU 2.0 on a 120 kV Talos L120C transmission electron microscope (Thermo Scientific) with a BM-Ceta camera. Data processing was done in CryoSPARC (Punjani et al. 2017), starting with CTF correction, particle picking and extraction. Three rounds of 2D classification were done, keeping only classes that had either HA monomer or trimer with bound Fabs. 3D models for these immune complexes were then generated using ab initio 3D reconstruction and heterogeneous refinement without imposing any symmetry. Classes that had clear, trimeric HA density and were representative of the diversity of Fab binding in each sample of polyclonal serum without redundancy were then separately subjected to a 3D refinement.

## Lead Contact

Further information and requests for resources and reagents should be directed to and will be fulfilled by the Lead Contact, Neil King.

## Acknowledgements

This work was funded by a generous gift from Open Philanthropy (N.P.K.); the Audacious Project at the Institute for Protein Design (N.P.K.); the Defense Threat Reduction Agency (HDTRA1-18-1-0001 to N.P.K.); the National Institute of Allergy and Infectious Diseases (P01 AI167966 to N.P.K.); and the intramural research program of the Vaccine Research Center, National Institute of Allergy and Infectious Diseases, National Institutes of Health (M.K.). This work was supported in part by the University of Washington’s Proteomics Resource (UWPR95794) and the UW Molecular Biophysics Training Program (T32GM008268). Labcorp performed all rabbit immunizations and blood draws.

## Author Contributions

A.D. and N.P.K. designed experiments and wrote the paper. A.D. and D.E. designed triheads. A.D. designed glycans and hypervariable triheads and carried out immunogen production and characterization, ELISAs, HAI, and ns-EMPEM analysis. S.B. and H.S. carried out neutralization assays. M.S. and K.K.L. designed and carried out HDX experiments. M.W. and M.G. performed glycan occupancy mass spectrometry. J.K. and M.N.P. handled all mouse immunization and blood draws.

## SUPPLEMENTARY FIGURES

**Figure S1.**
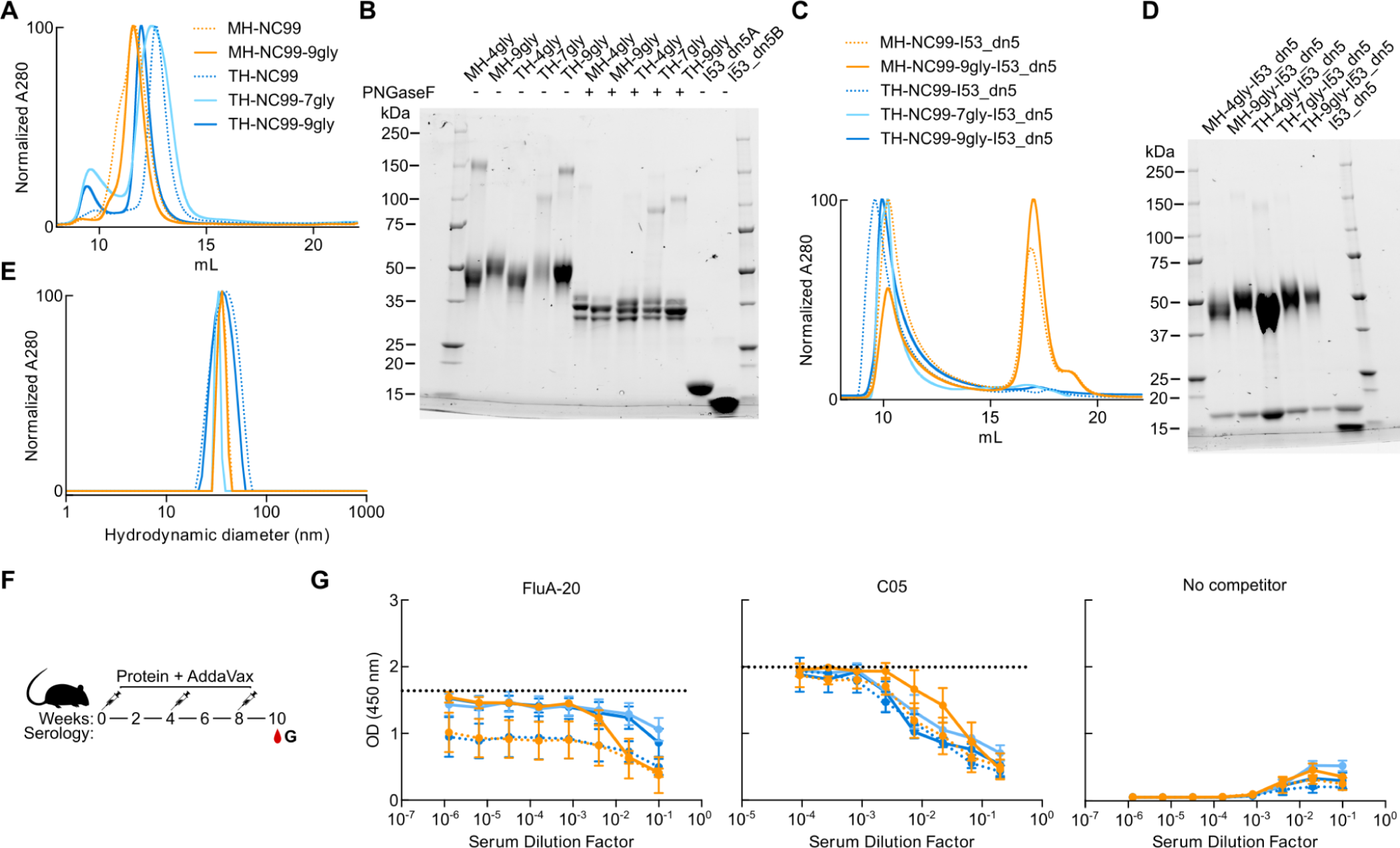
Hyperglycosylated TH-NC99-I53_dn5 Mouse Study. (A) SEC chromatograms of NC99 trihead nanoparticle components on a Superdex 200 Increase 10/300 GL column. (B) Reducing SDS-PAGE of wild-type and hyperglycosylated NC99 monoheads and trihead components without and with PNGaseF digestion, as well as bare I53_dn5B and I53_dn5A nanoparticle components. (C) NC99 trihead-I53_dn5 nanoparticle legend for Figures S1B, S1D, and S1F. SEC chromatograms of NC99 trihead-I53_dn5 nanoparticle immunogens on a Superose 6 Increase 10/300 GL column. (D) Reducing SDS-PAGE of NC99 trihead-I53_dn5 nanoparticles, as well as the bare I53_dn5 nanoparticle. (E) DLS of NC99 trihead-I53_dn5 nanoparticles. (F) Schematic illustrating mouse study timeline, immunizations, and serology timepoint. (G) Competition ELISA curves for NC99-foldon trimer ELISA antigen between week 10 sera and either FluA-20, C05, or a no competitor negative control. Dashed line is positive control of monoclonal binding in absence of sera.

**Figure S2.**
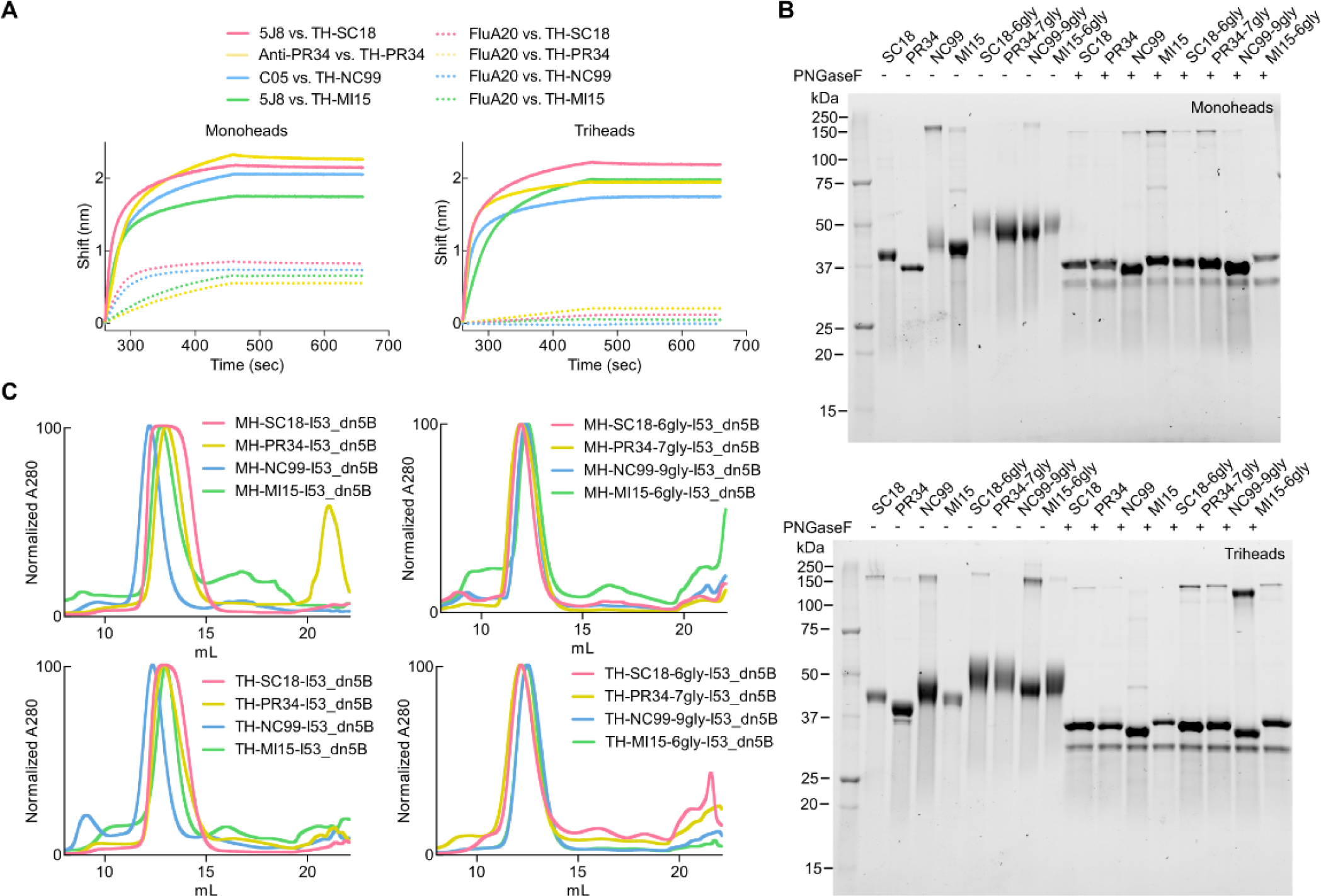
Mosaic and Hyperglycosylated Trihead Components Purification and Characterization. (A) BLI of monohead and trihead nanoparticle components against RBS-directed mAbs (5J8, anti-PR34, and C05) and FluA-20. (B) Reducing SDS-PAGE of wild-type and hyperglycosylated monohead and trihead nanoparticle components without and with PNGaseF digestion. (C) SEC chromatograms of wild-type and hyperglycosylated monohead and trihead nanoparticle components on a Superdex 200 Increase 10/300 GL column.

**Figure S3.**
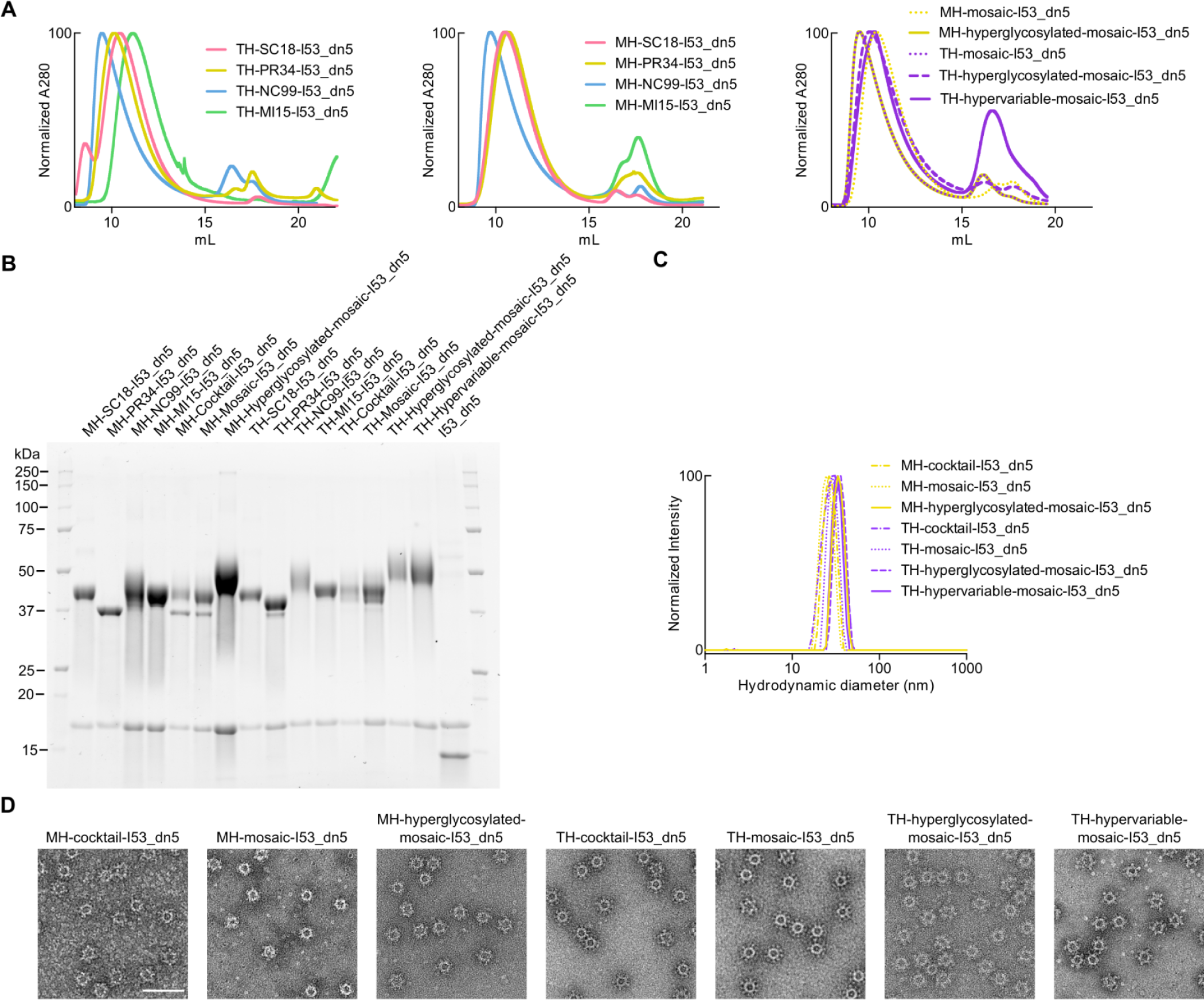
Mosaic, Hyperglycosylated, and Hypervariable Trihead Nanoparticles Purification and Characterization. (A) SEC chromatograms of individual H1 strains of monohead- and trihead-I53_dn5 nanoparticles and all mosaic nanoparticles on a Superose 6 Increase 10/300 GL column. (B) Reducing SDS-PAGE of individual H1 strains of monohead- and trihead-I53_dn5 nanoparticles and all mosaic nanoparticles, as well as the bare I53_dn5 nanoparticle. (C-D) C. DLS and D. nsEM of all mosaic nanoparticles. Scale bar = 100 nm.

**Figure S4.**
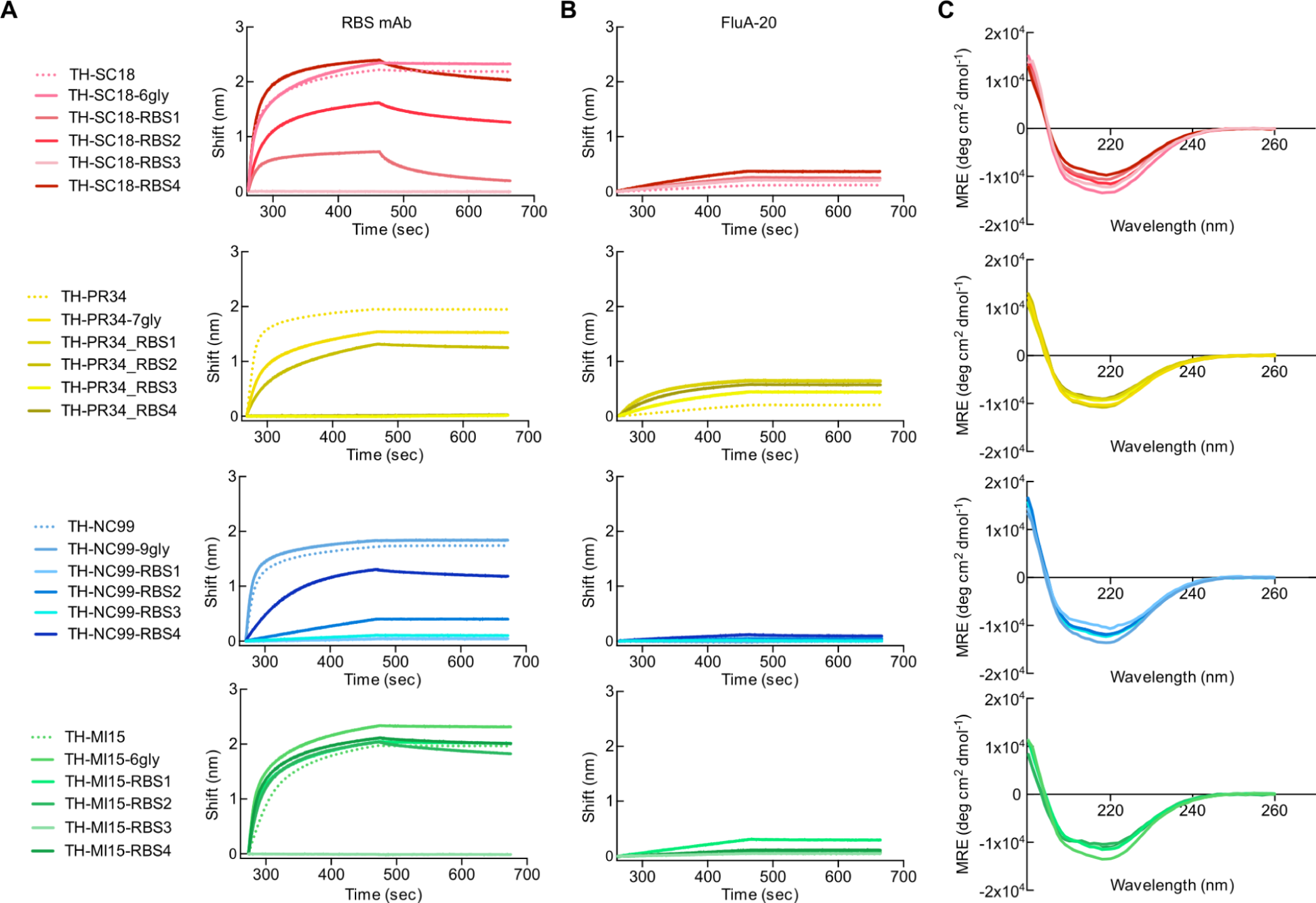
Hypervariable Trihead Immunogen Biophysical Characterization. (A) Legend of constructs in panels A-C. Binding of RBS-directed mAbs (5J8, anti-PR34, and C05) to all trihead components. (B) FluA-20 binding to all trihead components. (C) Far-UV circular dichroism (CD) spectra of all hypervariable trihead components.

**Figure S5.**
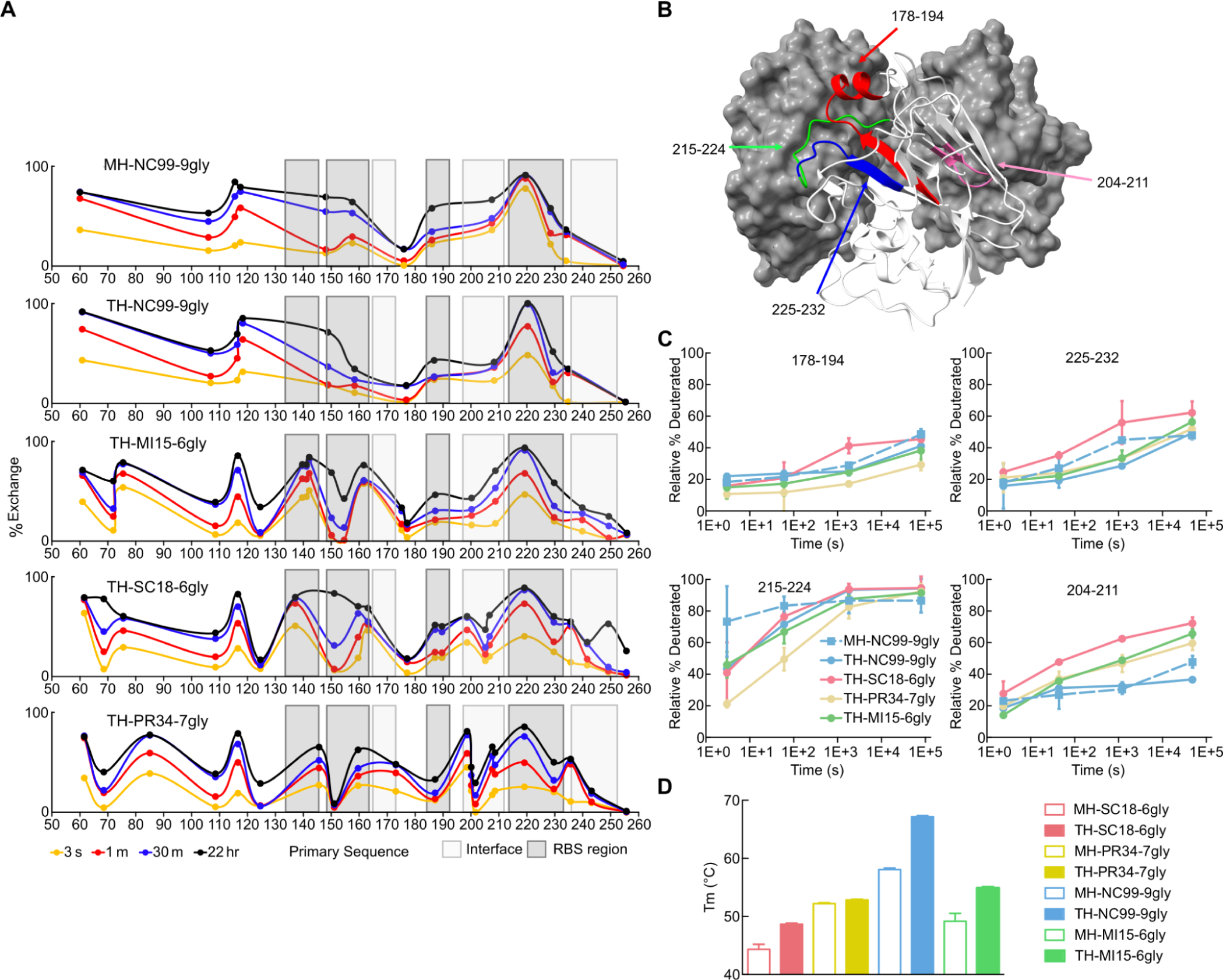
Hyperglycosylated Trihead Stability Characterization by HDX and Thermal Melts. (A) Deuterium uptake profiles across primary sequence plotted at four different timepoints for each construct. (B) HDX peptides in panel B highlighted by color on the SI06 (PDB: 5UG0) HA head domain. (C) Percent deuteration over time for two peptides in the RBS (178-194 and 225-232) and two peptides in the head trimer interface (215-224 and 204-211). (D) Melting temperatures of hyperglycosylated monoheads and triheads as measured by NanoDSF.

**Figure S6.**
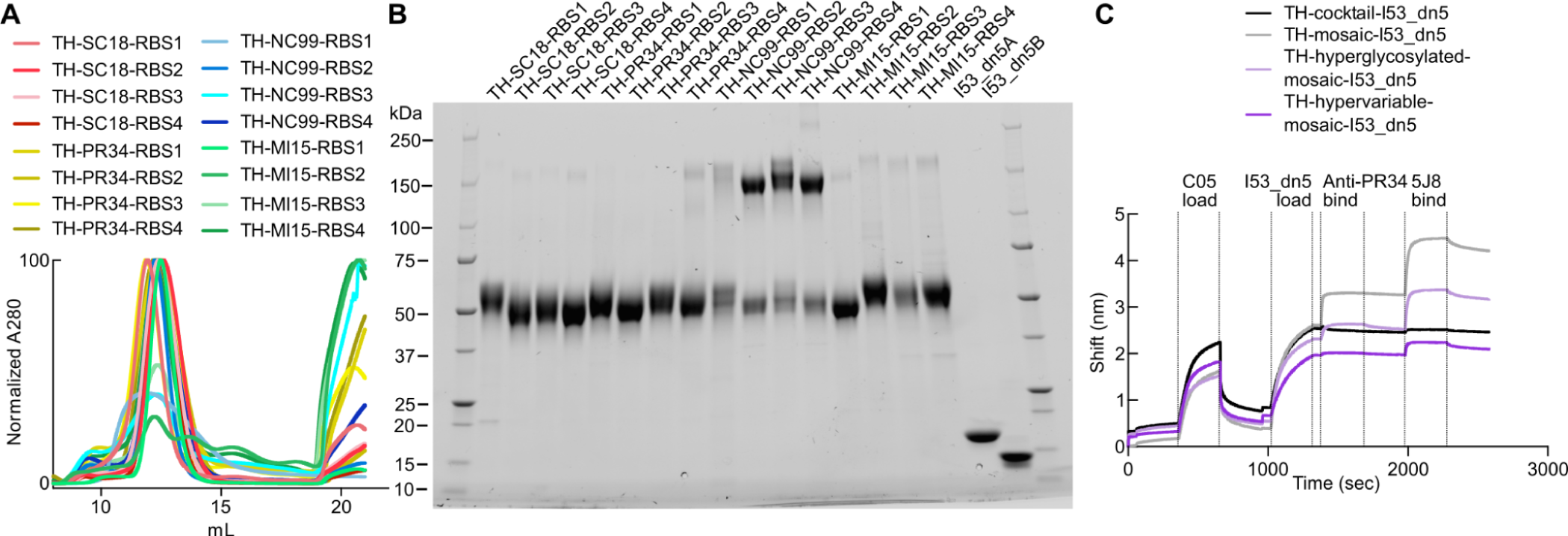
Hypervariable Trihead Immunogen Purification and Mosaic Nanoparticles BLI. (A) SEC chromatograms of hypervariable trihead nanoparticle components on a Superdex 200 Increase 10/300 GL column. (B) Reducing SDS-PAGE of hypervariable trihead-I53_dn5 nanoparticle components, as well as bare I53_dn5B and I53_dn5A nanoparticle components. (C) Sandwich BLI of trihead nanoparticle immunogens with C05 first captured on AR2G biosensors and then subsequent binding of nanoparticles, anti-PR34, and 5J8.

**Figure S7.**
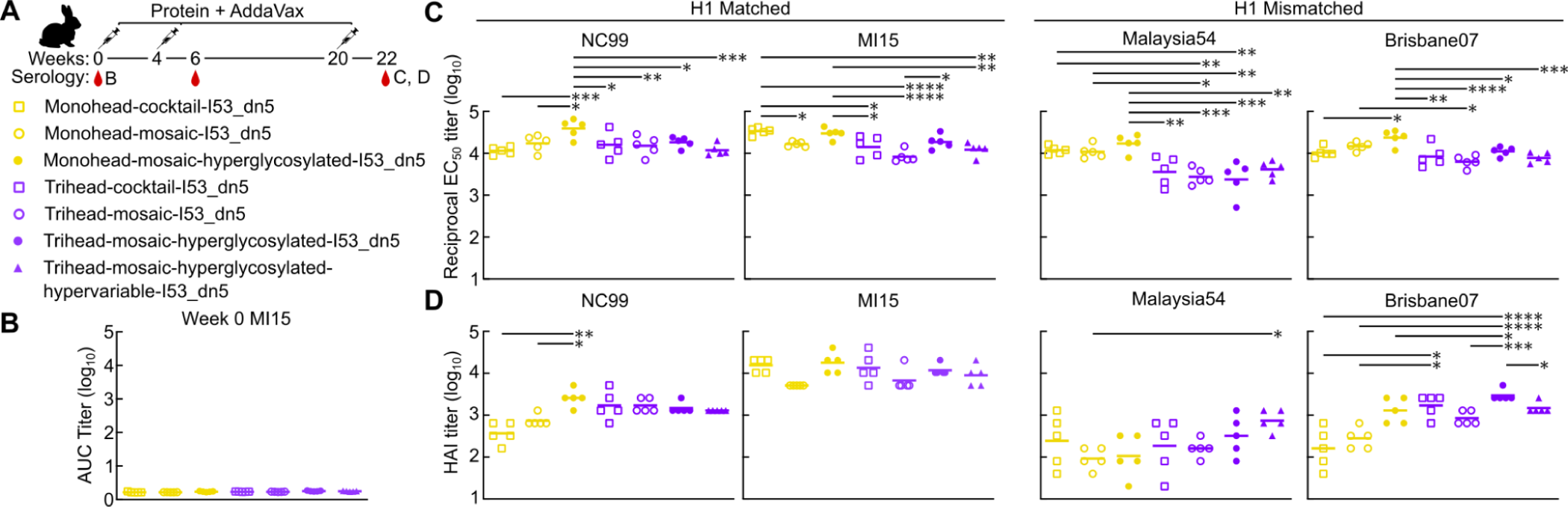
Vaccine-elicited Antibody Responses at Weeks 0 and 22 in Rabbits Immunized with Monohead and Trihead Nanoparticles. (A) Hypervariable trihead nanoparticle rabbit immunization schedule and groups. (B) ELISA AUC titers against vaccine-matched MI15 at week 0. (C-D) C. ELISA reciprocal EC_50_ titers and D. HAI titers at week 22, two weeks post second boost. Statistical significance was determined using one-way ANOVA with Tukey’s multiple comparisons test; ∗p < 0.05; ∗∗p < 0.01; ∗∗∗p < 0.001; ∗∗∗∗p < 0.0001.

**Table S1.**
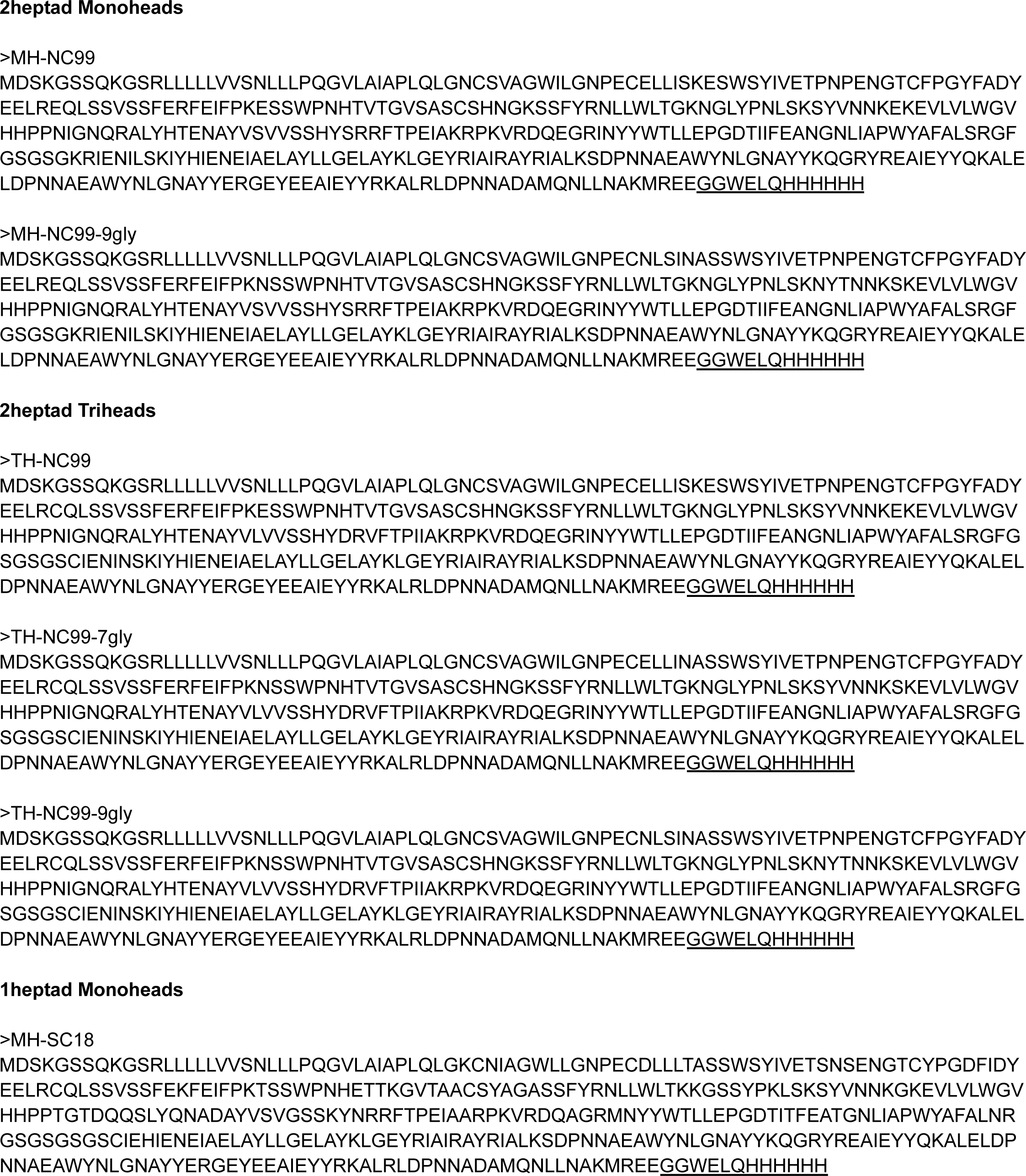
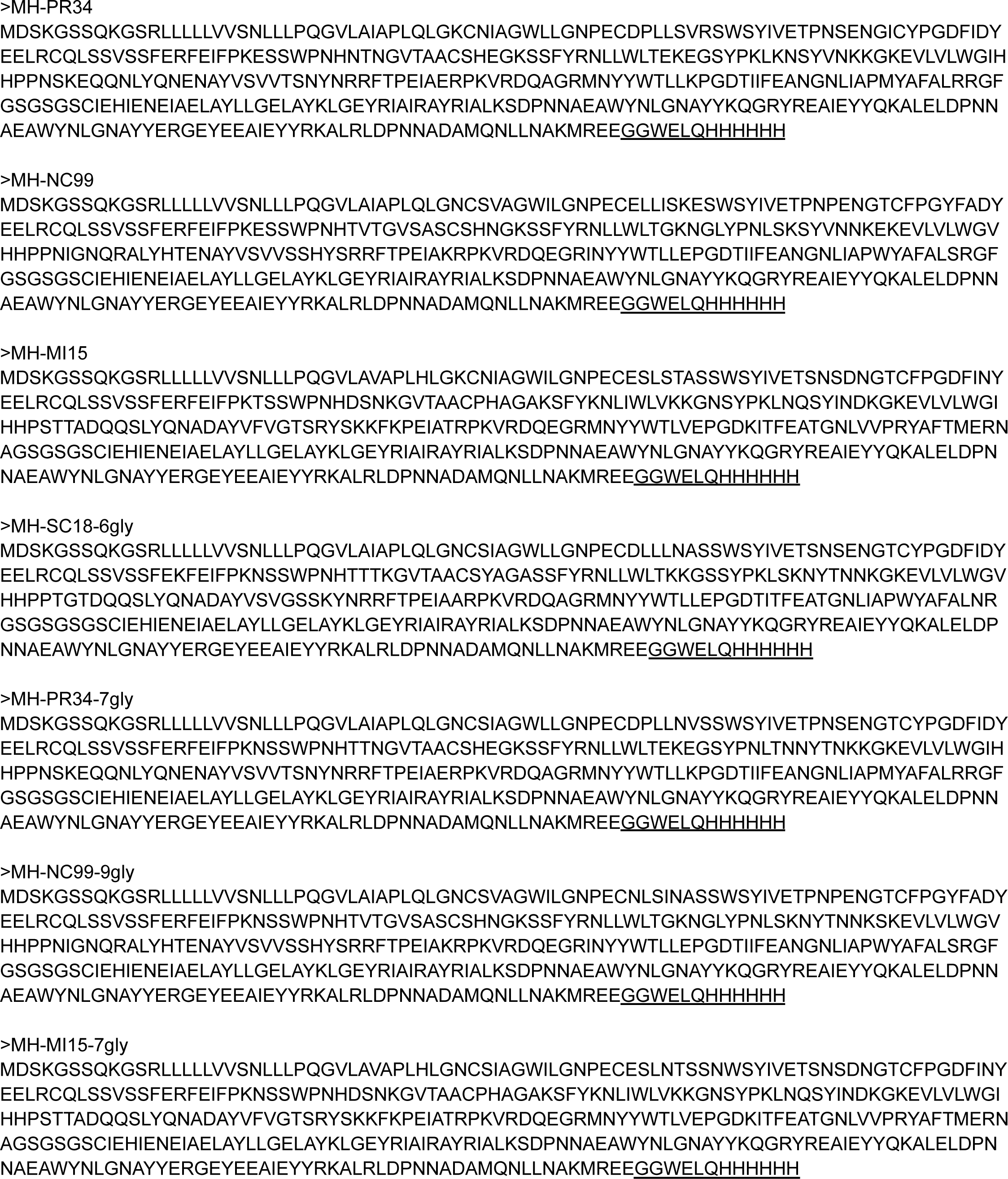
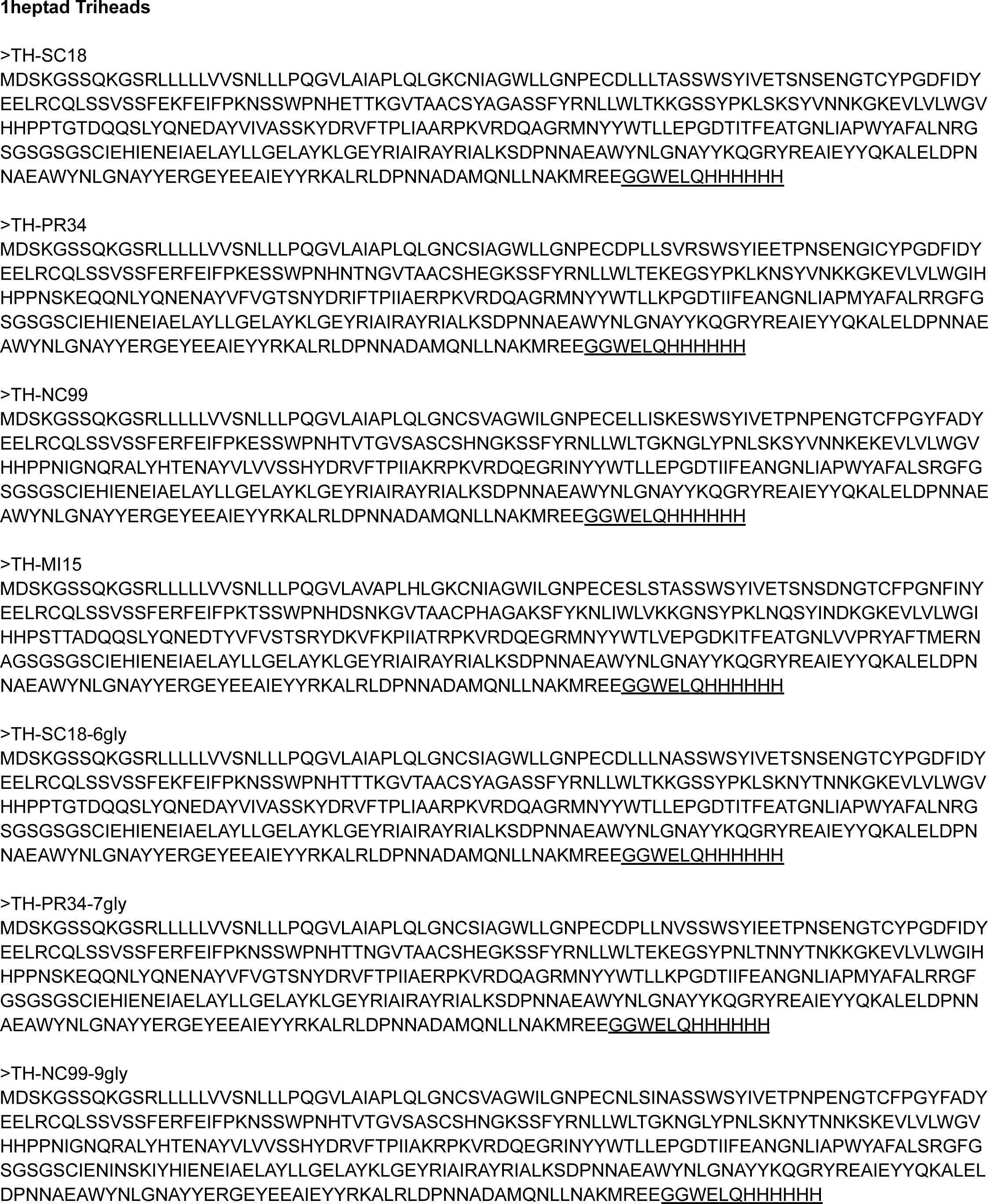
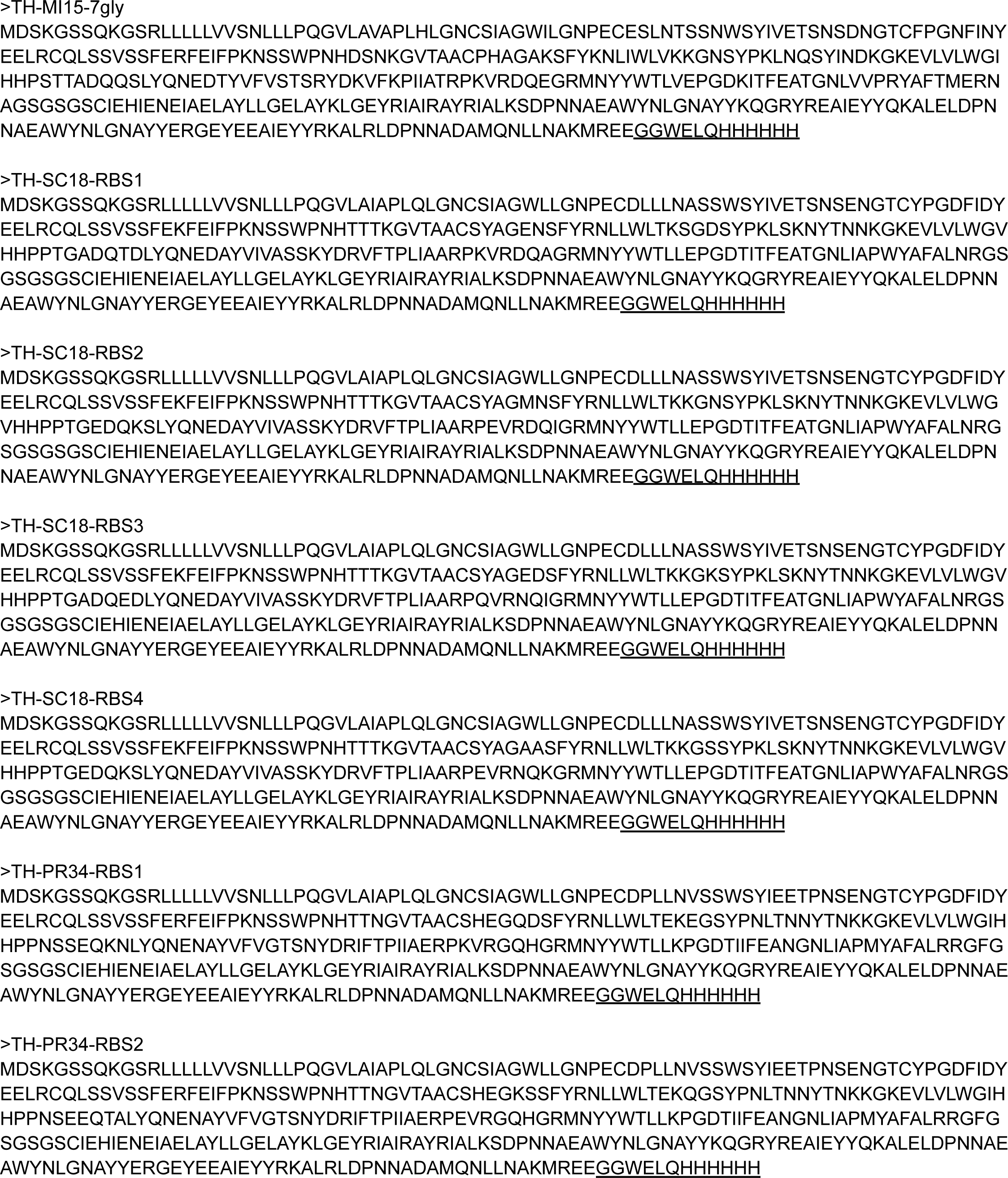
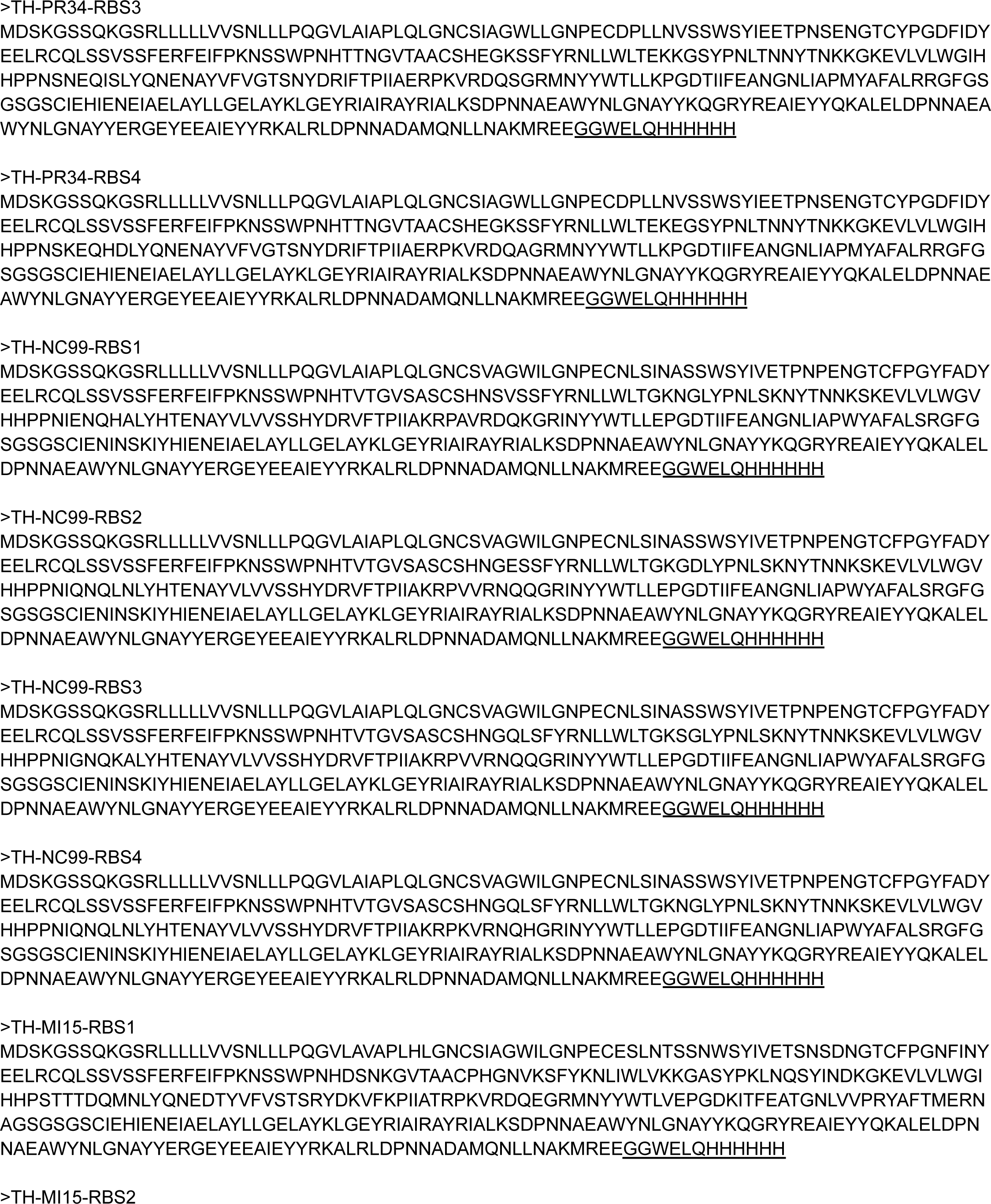
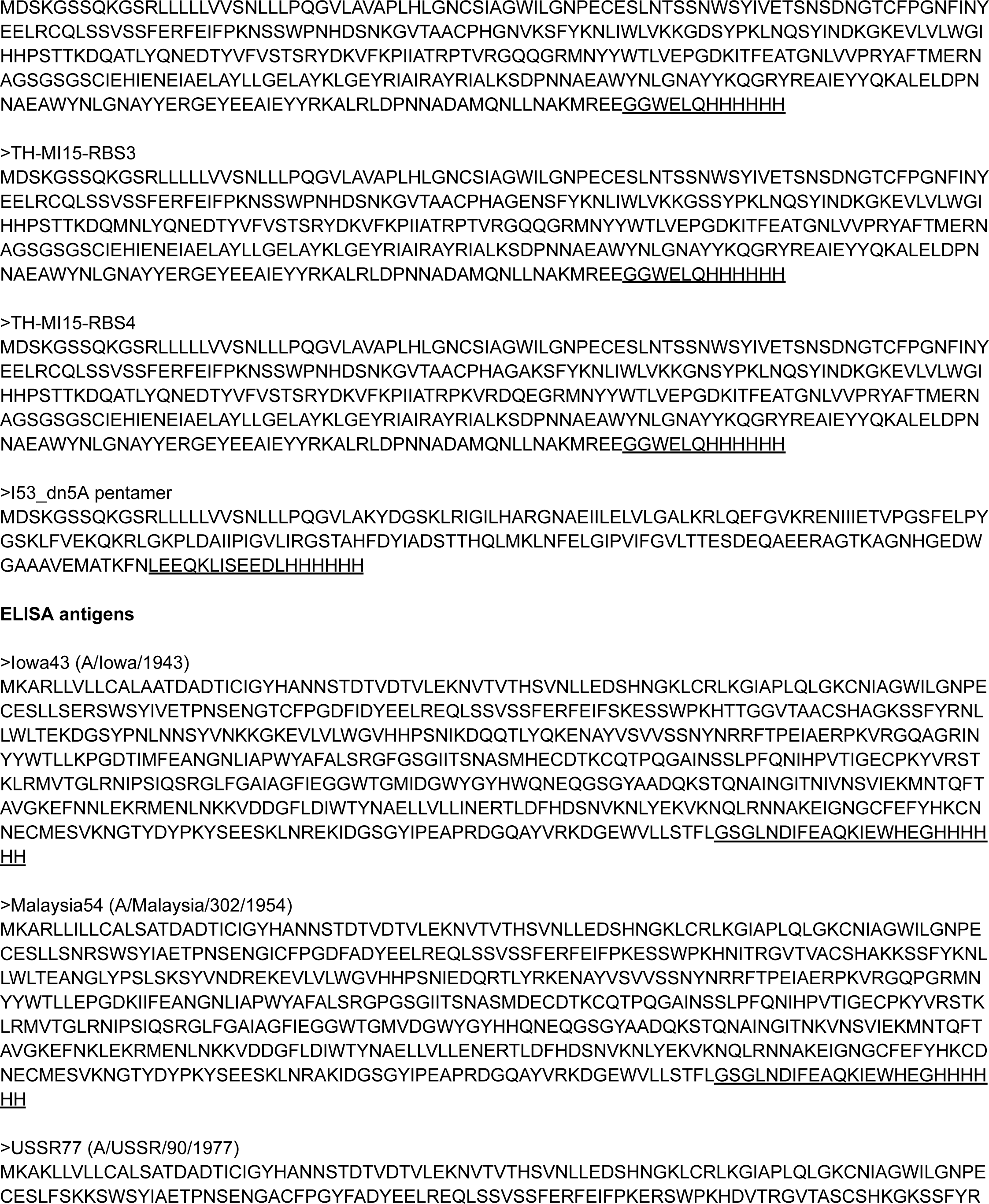
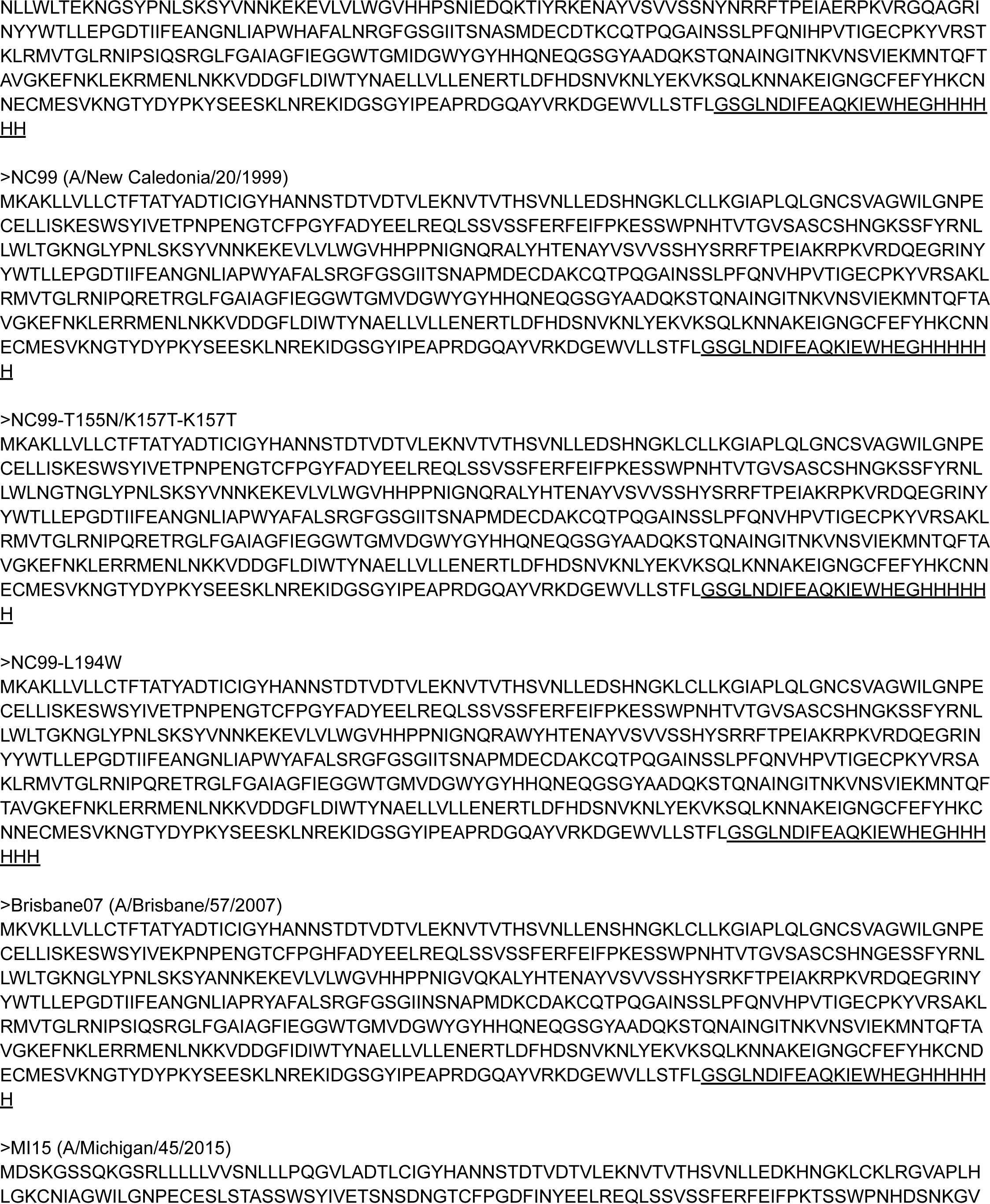
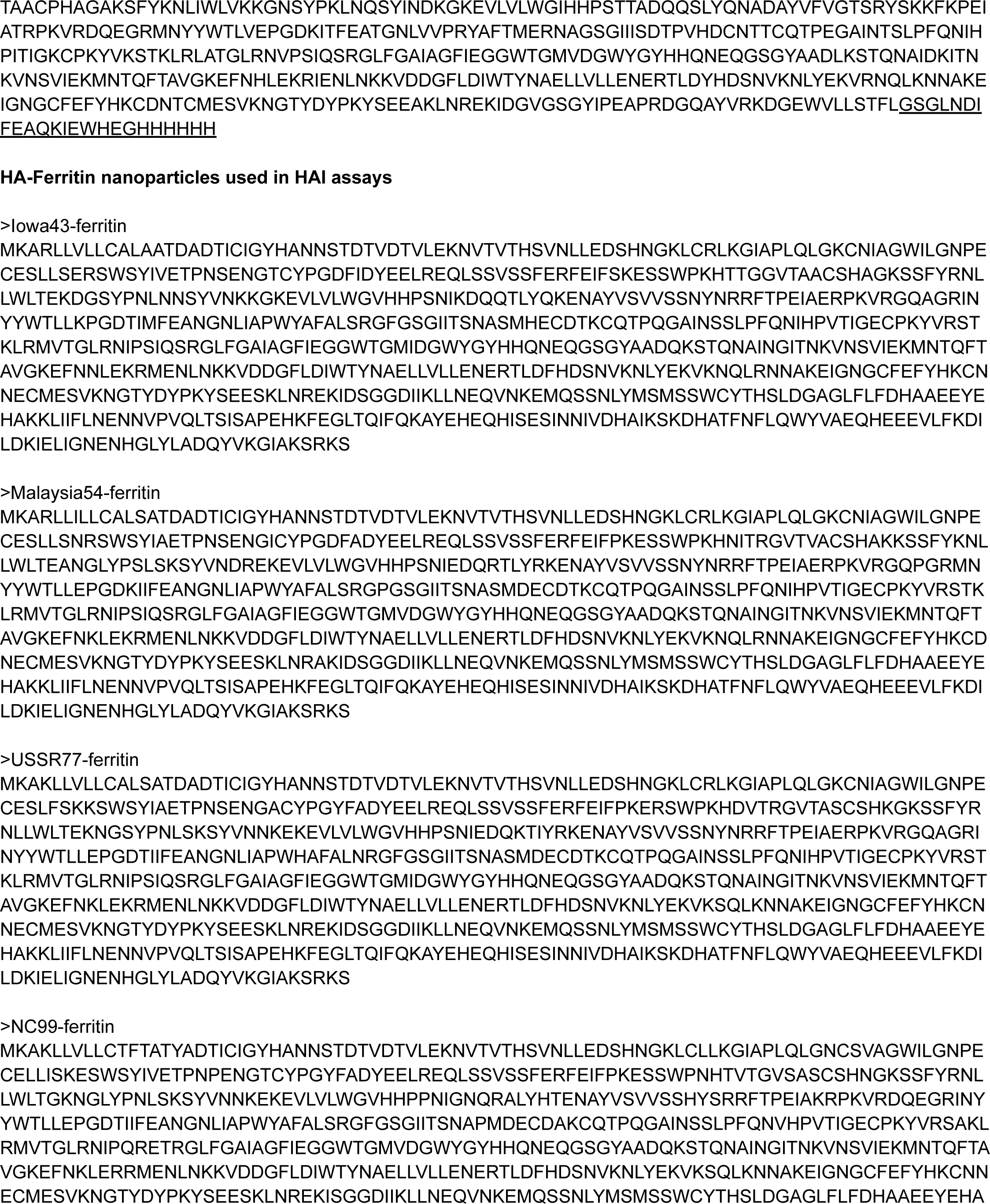
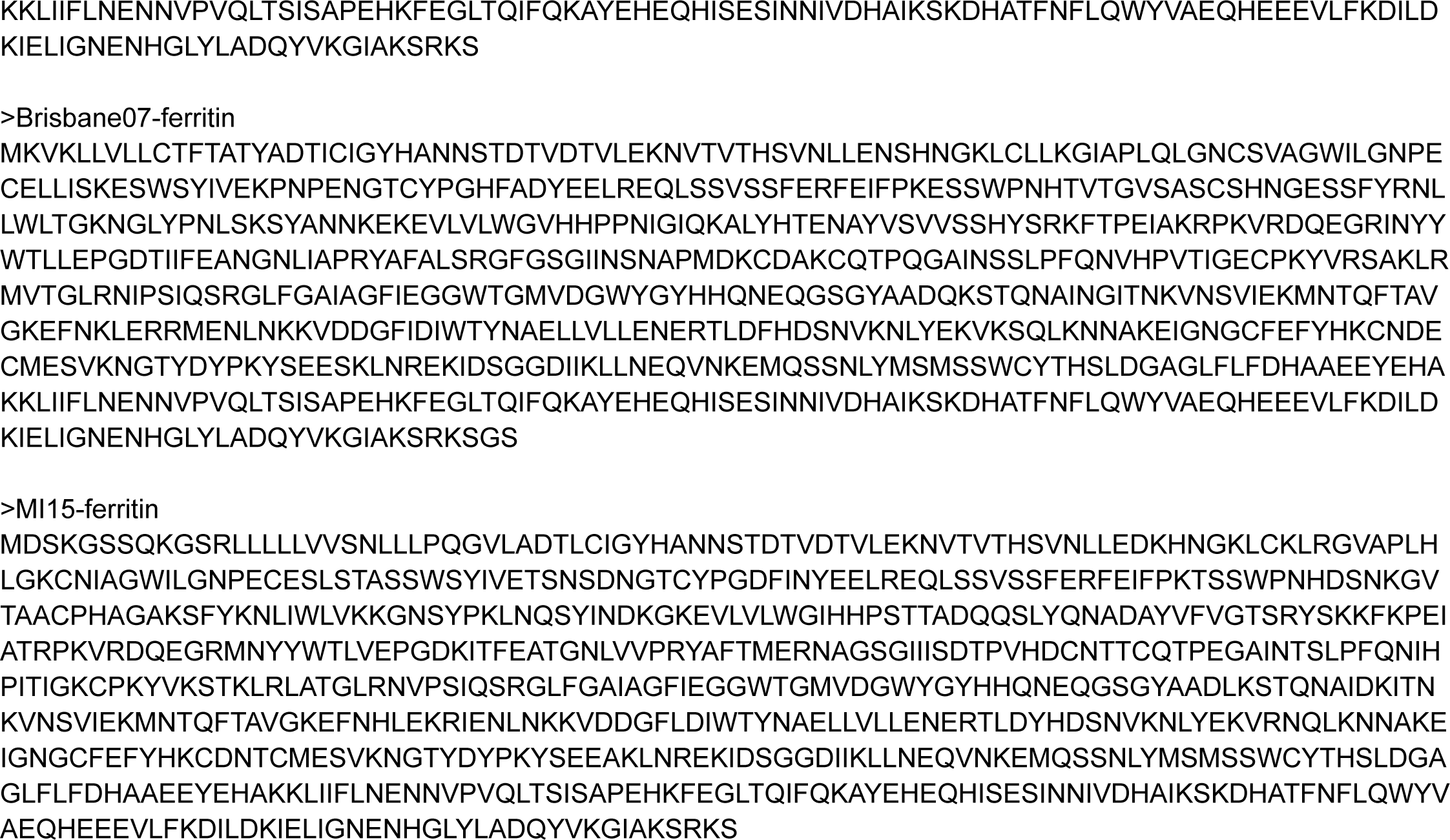
Amino acid sequences of novel proteins used in this study.

**Table S2.** Glycopeptide sequences and occupancies, related to Figure 1.

| Construct | Glycopeptide sequence | Sequon | Position | Occupancy (%) |
| --- | --- | --- | --- | --- |
| TH-NC99 | IAPLQLG**NCS**VAGWILGN<br>PE | G.NCS.V | 63 | 86.6 |
|  | **NGT**CFPGYFADYEE | E.NGT.C | 94 | 99 |
|  | ESSWP**NHT**VTGVSASCS<br>HNGK | P.NHT.V | 129 | 99.6 |
|  | NGLYP**NLS**K | P.NLS.K | 163 | 45.1 |
| TH-NC99-7gly | IAPLQLG**NCS**VAGWILGN<br>PE | G.NCS.V | 63 | 86.2 |
|  | LLIN**ASS**WSYIVET | I.NAS.S | 81 | 46.2 |
|  | **NGT**CFPGYFADYEE | E.NGT.C | 94 | 98.8 |
|  | **NSS**WP**NHT**VTGVSASCS<br>HNGK | K.NSS.W | 125b | 8.4 single,<br>91.5 double |
|  |  | P.NHT.V | 129 |  |
|  | NGLYP**NLS**K | P.NLS.K | 163 | 44.3 |
|  | SYV**NNK(S)** | N.NKS.K | 171 | 20.1 |
| TH-NC99-9gly | IAPLQLG**NCS**VAGWILGN<br>PE | G.NCS.V | 63 | 70 |
|  | **NLS**IN**ASS**WSYIVE | C.NLS.I | 77 | 63 single,<br>37 double |
|  |  | I.NAS.S | 81 |  |
|  | **NGT**CFPGYFADYEE | E.NGT.C | 94 | 99.6 |
|  | **NSS**WP**NHT**VTGVSASCS<br>HNGK | K.NSS.W | 125b | 5.0 single,<br>95.0 double |
|  |  | P.NHT.V | 129 |  |
|  | NGLYP**NLS**K | P.NLS.K | 163 | 44.2 |
|  | NGLYP**NLS**K**NYTNNK(S)** | P.NLS.K | 163 | 49.4 single,<br>40.8 double,<br>9.8 triple |
|  |  | K.NYT.N | 167 |  |
|  |  | N.NKS.K | 171 |  |

**Table S3.** Mutations introduced into hypervariable trihead antigens, related to Figure 3.

| Trihead RBS variant | Mutations |
| --- | --- |
| TH-SC18-RBS1 | A144E/S145N/K157S/S159D/T189A/Q192T/S193D |
| TH-SC18-RBS2 | A144M/S145N/S159N/T189E/Q192K/K222E/A227I |
| TH-SC18-RBS3 | A144E/S145D/S159K/T189A/Q192E/S193D/K222Q/<br>D225N/A227I |
| TH-SC18-RBS4 | S145A/T189E/Q192K/K222E/D225N/A227K |
| TH-PR34-RBS1 | K144Q/S145D/K189S/Q192K/D225G/A227H |
| TH-PR34-RBS2 | E158Q/K189E/Q192T/N193A/K222E/D225G/A227H |
| TH-PR34-RBS3 | E158K/K189N/Q192I/N193S/A227S |
| TH-PR34-RBS4 | Q192H/N193D |
| TH-NC99-RBS1 | G143S/K144V/G189E/R192H/K222A/E227K |
| TH-NC99-RBS2 | K144E/N158G/G159D/G189Q/R192L/A193N/K222V/<br>E227Q |
| TH-NC99-RBS3 | K144Q/S145L/N158S/R192K/K222V/D225N/E227Q |
| TH-NC99-RBS4 | K144Q/S145L/G189Q/R192L/A193N/D225N/E227H |
| TH-MI15-RBS1 | A142G/G143N/A144V/A189T/Q192M/S193N |
| TH-MI15-RBS2 | A142G/G143N/A144V/N159D/A189K/Q192A/S193T/<br>K222T/Q225G/G227Q |
| TH-MI15-RBS3 | A144E/K145N/N159S/A189K/Q192M/S193N/K222T/<br>Q225G/G227Q |
| TH-MI15-RBS4 | A189K/Q192A/S193T |

## References

Abbott, Robert K., Jeong Hyun Lee, Sergey Menis, Patrick Skog, Meghan Rossi, Takayuki Ota, Daniel W. Kulp, et al. 2018. “Precursor Frequency and Affinity Determine B Cell Competitive Fitness in Germinal Centers, Tested with Germline-Targeting HIV Vaccine Immunogens.” Immunity 48 (1): 133–46.e6.

Adolf-Bryfogle, Jared, Jason W. Labonte, John C. Kraft, Maxim Shapovalov, Sebastian Raemisch, Thomas Lütteke, Frank DiMaio, et al. 2021. “Growing Glycans in Rosetta: Accurate de Novo Glycan Modeling, Density Fitting, and Rational Sequon Design.” bioRxiv. https://doi.org/10.1101/2021.09.27.462000.

Altman, Meghan O., Davide Angeletti, and Jonathan W. Yewdell. 2018. “Antibody Immunodominance: The Key to Understanding Influenza Virus Antigenic Drift.” Viral Immunology 31 (2): 142–49.

Altman, Meghan O., Jack R. Bennink, Jonathan W. Yewdell, and Brantley R. Herrin. 2015. “Lamprey VLRB Response to Influenza Virus Supports Universal Rules of Immunogenicity and Antigenicity.” eLife 4 (August). https://doi.org/10.7554/eLife.07467.

Andrews, Sarah F., Lauren Y. Cominsky, Geoffrey D. Shimberg, Rebecca A. Gillespie, Jason Gorman, Julie E. Raab, Joshua Brand, et al. 2023. “An Influenza H1 Hemagglutinin Stem-Only Immunogen Elicits a Broadly Cross-Reactive B Cell Response in Humans.” Science Translational Medicine 15 (692): eade4976.

Angeletti, Davide, James S. Gibbs, Matthew Angel, Ivan Kosik, Heather D. Hickman, Gregory M. Frank, Suman R. Das, et al. 2017. “Defining B Cell Immunodominance to Viruses.” Nature Immunology 18 (4): 456–63.

Aung, Aereas, Ang Cui, Laura Maiorino, Ava P. Amini, Justin R. Gregory, Maurice Bukenya, Yiming Zhang, et al. 2023. “Low Protease Activity in B Cell Follicles Promotes Retention of Intact Antigens after Immunization.” Science 379 (6630): eabn8934.

Bajic, Goran, Max J. Maron, Yu Adachi, Taishi Onodera, Kevin R. McCarthy, Charles E. McGee, Gregory D. Sempowski, et al. 2019. “Influenza Antigen Engineering Focuses Immune Responses to a Subdominant but Broadly Protective Viral Epitope.” Cell Host & Microbe 25 (6): 827–35.e6.

Bangaru, Sandhya, Shanshan Lang, Michael Schotsaert, Hillary A. Vanderven, Xueyong Zhu, Nurgun Kose, Robin Bombardi, et al. 2019. “A Site of Vulnerability on the Influenza Virus Hemagglutinin Head Domain Trimer Interface.” Cell 177 (5): 1136–52.e18.

Barouch, Dan H., Zhi-Yong Yang, Wing-Pui Kong, Birgit Korioth-Schmitz, Shawn M. Sumida, Diana M. Truitt, Michael G. Kishko, et al. 2005. “A Human T-Cell Leukemia Virus Type 1 Regulatory Element Enhances the Immunogenicity of Human Immunodeficiency Virus Type 1 DNA Vaccines in Mice and Nonhuman Primates.” Journal of Virology 79 (14): 8828–34.

Bedford, Trevor, Marc A. Suchard, Philippe Lemey, Gytis Dudas, Victoria Gregory, Alan J. Hay, John W. McCauley, Colin A. Russell, Derek J. Smith, and Andrew Rambaut. 2014. “Integrating Influenza Antigenic Dynamics with Molecular Evolution.” eLife 3 (February): e01914.

Benton, Donald J., Andrea Nans, Lesley J. Calder, Jack Turner, Ursula Neu, Yi Pu Lin, Esther Ketelaars, et al. 2018. “Influenza Hemagglutinin Membrane Anchor.” Proceedings of the National Academy of Sciences of the United States of America 115 (40): 10112–17.

Bianchi, Matteo, Hannah L. Turner, Bartek Nogal, Christopher A. Cottrell, David Oyen, Matthias Pauthner, Raiza Bastidas, et al. 2018. “Electron-Microscopy-Based Epitope Mapping Defines Specificities of Polyclonal Antibodies Elicited during HIV-1 BG505 Envelope Trimer Immunization.” Immunity 49 (2): 288–300.e8.

Boyoglu-Barnum, Seyhan, Daniel Ellis, Rebecca A. Gillespie, Geoffrey B. Hutchinson, Young-Jun Park, Syed M. Moin, Oliver J. Acton, et al. 2021. “Quadrivalent Influenza Nanoparticle Vaccines Induce Broad Protection.” Nature 592 (7855): 623–28.

Brinkkemper, Mitch, Tim S. Veth, Philip J. M. Brouwer, Hannah Turner, Meliawati Poniman, Judith A. Burger, Joey H. Bouhuijs, et al. 2022. “Co-Display of Diverse Spike Proteins on Nanoparticles Broadens Sarbecovirus Neutralizing Antibody Responses.” iScience 25 (12): 105649.

Broecker, Felix, Sean T. H. Liu, Nungruthai Suntronwong, Weina Sun, Mark J. Bailey, Raffael Nachbagauer, Florian Krammer, and Peter Palese. 2019. “A Mosaic Hemagglutinin-Based Influenza Virus Vaccine Candidate Protects Mice from Challenge with Divergent H3N2 Strains.” NPJ Vaccines 4 (July): 31.

Carrat, F., and A. Flahault. 2007. “Influenza Vaccine: The Challenge of Antigenic Drift.” Vaccine 25 (39-40): 6852–62.

Cirelli, Kimberly M., Diane G. Carnathan, Bartek Nogal, Jacob T. Martin, Oscar L. Rodriguez, Amit A. Upadhyay, Chiamaka A. Enemuo, et al. 2019. “Slow Delivery Immunization Enhances HIV Neutralizing Antibody and Germinal Center Responses via Modulation of Immunodominance.” Cell 177 (5): 1153–71.e28.

Cohen, Alexander A., Neeltje van Doremalen, Allison J. Greaney, Hanne Andersen, Ankur Sharma, Tyler N. Starr, Jennifer R. Keeffe, et al. 2022. “Mosaic RBD Nanoparticles Protect against Challenge by Diverse Sarbecoviruses in Animal Models.” Science 377 (6606): eabq0839.

Cohen, Alexander A., Priyanthi N. P. Gnanapragasam, Yu E. Lee, Pauline R. Hoffman, Susan Ou, Leesa M. Kakutani, Jennifer R. Keeffe, et al. 2021. “Mosaic Nanoparticles Elicit Cross-Reactive Immune Responses to Zoonotic Coronaviruses in Mice.” Science 371 (6530): 735–41.

Cohen, Alexander A., Zhi Yang, Priyanthi N. P. Gnanapragasam, Susan Ou, Kim-Marie A. Dam, Haoqing Wang, and Pamela J. Bjorkman. 2021. “Construction, Characterization, and Immunization of Nanoparticles That Display a Diverse Array of Influenza HA Trimers.” PloS One 16 (3): e0247963.

Corbett, Kizzmekia S., Syed M. Moin, Hadi M. Yassine, Alberto Cagigi, Masaru Kanekiyo, Seyhan Boyoglu-Barnum, Sky I. Myers, et al. 2019. “Design of Nanoparticulate Group 2 Influenza Virus Hemagglutinin Stem Antigens That Activate Unmutated Ancestor B Cell Receptors of Broadly Neutralizing Antibody Lineages.” mBio 10 (1). https://doi.org/10.1128/mBio.02810-18.

Creanga, Adrian, Rebecca A. Gillespie, Brian E. Fisher, Sarah F. Andrews, Julia Lederhofer, Christina Yap, Liam Hatch, et al. 2021. “A Comprehensive Influenza Reporter Virus Panel for High-Throughput Deep Profiling of Neutralizing Antibodies.” Nature Communications 12 (1): 1722.

Darricarrère, Nicole, Yu Qiu, Masaru Kanekiyo, Adrian Creanga, Rebecca A. Gillespie, Syed M. Moin, Jacqueline Saleh, et al. 2021. “Broad Neutralization of H1 and H3 Viruses by Adjuvanted Influenza HA Stem Vaccines in Nonhuman Primates.” Science Translational Medicine 13 (583). https://doi.org/10.1126/scitranslmed.abe5449.

Doud, Michael B., and Jesse D. Bloom. 2016. “Accurate Measurement of the Effects of All Amino-Acid Mutations on Influenza Hemagglutinin.” Viruses 8 (6). https://doi.org/10.3390/v8060155.

Dreyfus, Cyrille, Nick S. Laursen, Ted Kwaks, David Zuijdgeest, Reza Khayat, Damian C. Ekiert, Jeong Hyun Lee, et al. 2012. “Highly Conserved Protective Epitopes on Influenza B Viruses.” Science 337 (6100): 1343–48.

Duan, Hongying, Xuejun Chen, Jeffrey C. Boyington, Cheng Cheng, Yi Zhang, Alexander J. Jafari, Tyler Stephens, et al. 2018. “Glycan Masking Focuses Immune Responses to the HIV-1 CD4-Binding Site and Enhances Elicitation of VRC01-Class Precursor Antibodies.” Immunity 49 (2): 301–11.e5.

Eggink, Dirk, Peter H. Goff, and Peter Palese. 2014. “Guiding the Immune Response against Influenza Virus Hemagglutinin toward the Conserved Stalk Domain by Hyperglycosylation of the Globular Head Domain.” Journal of Virology 88 (1): 699–704.

Ekiert, Damian C., Arun K. Kashyap, John Steel, Adam Rubrum, Gira Bhabha, Reza Khayat, Jeong Hyun Lee, et al. 2012. “Cross-Neutralization of Influenza A Viruses Mediated by a Single Antibody Loop.” Nature 489 (7417): 526–32.

Ellis, Daniel, Annie Dosey, Seyhan Boyoglu-Barnum, Young-Jun Park, Rebecca Gillespie, Hubza Syeda, Yaroslav Tsybovsky, et al. n.d. “Antigen-Antigen Spacing on Protein Nanoparticles Influences Antibody Responses to Vaccination.”

Ellis, Daniel, Julia Lederhofer, Oliver J. Acton, Yaroslav Tsybovsky, Sally Kephart, Christina Yap, Rebecca A. Gillespie, et al. 2022. “Structure-Based Design of Stabilized Recombinant Influenza Neuraminidase Tetramers.” Nature Communications 13 (1): 1825.

Guthmiller, Jenna J., Julianna Han, Henry A. Utset, Lei Li, Linda Yu-Ling Lan, Carole Henry, Christopher T. Stamper, et al. 2022. “Broadly Neutralizing Antibodies Target a Haemagglutinin Anchor Epitope.” Nature 602 (7896): 314–20.

Han, Julianna, Aaron J. Schmitz, Sara T. Richey, Ya-Nan Dai, Hannah L. Turner, Bassem M. Mohammed, Daved H. Fremont, Ali H. Ellebedy, and Andrew B. Ward. 2021. “Polyclonal Epitope Mapping Reveals Temporal Dynamics and Diversity of Human Antibody Responses to H5N1 Vaccination.” Cell Reports 34 (4): 108682.

Hsieh, Ching-Lin, Jory A. Goldsmith, Jeffrey M. Schaub, Andrea M. DiVenere, Hung-Che Kuo, Kamyab Javanmardi, Kevin C. Le, et al. 2020. “Structure-Based Design of Prefusion-Stabilized SARS-CoV-2 Spikes.” Science 369 (6510): 1501–5.

Kallewaard, Nicole L., Davide Corti, Patrick J. Collins, Ursula Neu, Josephine M. McAuliffe, Ebony Benjamin, Leslie Wachter-Rosati, et al. 2016. “Structure and Function Analysis of an Antibody Recognizing All Influenza A Subtypes.” Cell 166 (3): 596–608.

Kanekiyo, Masaru, M. Gordon Joyce, Rebecca A. Gillespie, John R. Gallagher, Sarah F. Andrews, Hadi M. Yassine, Adam K. Wheatley, et al. 2019. “Mosaic Nanoparticle Display of Diverse Influenza Virus Hemagglutinins Elicits Broad B Cell Responses.” Nature Immunology 20 (3): 362–72.

Kanekiyo, Masaru, Chih-Jen Wei, Hadi M. Yassine, Patrick M. McTamney, Jeffrey C. Boyington, James R. R. Whittle, Srinivas S. Rao, Wing-Pui Kong, Lingshu Wang, and Gary J. Nabel. 2013. “Self-Assembling Influenza Nanoparticle Vaccines Elicit Broadly Neutralizing H1N1 Antibodies.” Nature 499 (7456): 102–6.

Kato, Yu, Robert K. Abbott, Brian L. Freeman, Sonya Haupt, Bettina Groschel, Murillo Silva, Sergey Menis, Darrell J. Irvine, William R. Schief, and Shane Crotty. 2020. “Multifaceted Effects of Antigen Valency on B Cell Response Composition and Differentiation In Vivo.” Immunity 53 (3): 548–63.e8.

Kobayashi, Yuki, and Yoshiyuki Suzuki. 2012. “Evidence for N-Glycan Shielding of Antigenic Sites during Evolution of Human Influenza A Virus Hemagglutinin.” Journal of Virology 86 (7): 3446–51.

Koel, Björn F., David F. Burke, Theo M. Bestebroer, Stefan van der Vliet, Gerben C. M. Zondag, Gaby Vervaet, Eugene Skepner, et al. 2013. “Substitutions near the Receptor Binding Site Determine Major Antigenic Change during Influenza Virus Evolution.” Science 342 (6161): 976–79.

Krarup, Anders, Daphné Truan, Polina Furmanova-Hollenstein, Lies Bogaert, Pascale Bouchier, Ilona J. M. Bisschop, Myra N. Widjojoatmodjo, et al. 2015. “A Highly Stable Prefusion RSV F Vaccine Derived from Structural Analysis of the Fusion Mechanism.” Nature Communications 6 (September): 8143.

Krause, Jens C., Tshidi Tsibane, Terrence M. Tumpey, Chelsey J. Huffman, Christopher F. Basler, and James E. Crowe Jr. 2011. “A Broadly Neutralizing Human Monoclonal Antibody That Recognizes a Conserved, Novel Epitope on the Globular Head of the Influenza H1N1 Virus Hemagglutinin.” Journal of Virology 85 (20): 10905–8.

Lee, Jiwon, Daniel R. Boutz, Veronika Chromikova, M. Gordon Joyce, Christopher Vollmers, Kwanyee Leung, Andrew P. Horton, et al. 2016. “Molecular-Level Analysis of the Serum Antibody Repertoire in Young Adults before and after Seasonal Influenza Vaccination.” Nature Medicine 22 (12): 1456–64.

Lee, Peter S., Reiko Yoshida, Damian C. Ekiert, Naoki Sakai, Yasuhiko Suzuki, Ayato Takada, and Ian A. Wilson. 2012. “Heterosubtypic Antibody Recognition of the Influenza Virus Hemagglutinin Receptor Binding Site Enhanced by Avidity.” Proceedings of the National Academy of Sciences of the United States of America 109 (42): 17040–45.

Leman, Julia Koehler, Brian D. Weitzner, Steven M. Lewis, Jared Adolf-Bryfogle, Nawsad Alam, Rebecca F. Alford, Melanie Aprahamian, et al. 2020. “Macromolecular Modeling and Design in Rosetta: Recent Methods and Frameworks.” Nature Methods 17 (7): 665–80.

Li, Tingting, Junyu Chen, Qingbing Zheng, Wenhui Xue, Limin Zhang, Rui Rong, Sibo Zhang, et al. 2022. “Identification of a Cross-Neutralizing Antibody That Targets the Receptor Binding Site of H1N1 and H5N1 Influenza Viruses.” Nature Communications 13 (1): 5182.

McCarthy, Kevin R., Akiko Watanabe, Masayuki Kuraoka, Khoi T. Do, Charles E. McGee, Gregory D. Sempowski, Thomas B. Kepler, Aaron G. Schmidt, Garnett Kelsoe, and Stephen C. Harrison. 2018. “Memory B Cells That Cross-React with Group 1 and Group 2 Influenza A Viruses Are Abundant in Adult Human Repertoires.” Immunity 48 (1): 174–84.e9.

Moin, Syed M., Jeffrey C. Boyington, Seyhan Boyoglu-Barnum, Rebecca A. Gillespie, Gabriele Cerutti, Crystal Sao-Fong Cheung, Alberto Cagigi, et al. 2022. “Co-Immunization with Hemagglutinin Stem Immunogens Elicits Cross-Group Neutralizing Antibodies and Broad Protection against Influenza A Viruses.” *Immunity*, November. https://doi.org/10.1016/j.immuni.2022.10.015.

Pallesen, Jesper, Nianshuang Wang, Kizzmekia S. Corbett, Daniel Wrapp, Robert N. Kirchdoerfer, Hannah L. Turner, Christopher A. Cottrell, et al. 2017. “Immunogenicity and Structures of a Rationally Designed Prefusion MERS-CoV Spike Antigen.” Proceedings of the National Academy of Sciences of the United States of America 114 (35): E7348–57.

Pino, Lindsay K., Brian C. Searle, James G. Bollinger, Brook Nunn, Brendan MacLean, and Michael J. MacCoss. 2020. “The Skyline Ecosystem: Informatics for Quantitative Mass Spectrometry Proteomics.” Mass Spectrometry Reviews 39 (3): 229–44.

Punjani, Ali, John L. Rubinstein, David J. Fleet, and Marcus A. Brubaker. 2017. “cryoSPARC: Algorithms for Rapid Unsupervised Cryo-EM Structure Determination.” Nature Methods 14 (3): 290–96.

Raymond, Donald D., Goran Bajic, Jack Ferdman, Pirada Suphaphiphat, Ethan C. Settembre, M. Anthony Moody, Aaron G. Schmidt, and Stephen C. Harrison. 2018. “Conserved Epitope on Influenza-Virus Hemagglutinin Head Defined by a Vaccine-Induced Antibody.” Proceedings of the National Academy of Sciences 115 (1): 168–73.

Read, Benjamin J., Lori Won, John C. Kraft, Isaac Sappington, Aereas Aung, Shengwei Wu, Julia Bals, et al. 2022. “Mannose-Binding Lectin and Complement Mediate Follicular Localization and Enhanced Immunogenicity of Diverse Protein Nanoparticle Immunogens.” Cell Reports 38 (2): 110217.

Rejmanek, Daniel, Parviez R. Hosseini, Jonna A. K. Mazet, Peter Daszak, and Tracey Goldstein. 2015. “Evolutionary Dynamics and Global Diversity of Influenza A Virus.” Journal of Virology 89 (21): 10993–1.

Sanders, Rogier W., Ronald Derking, Albert Cupo, Jean-Philippe Julien, Anila Yasmeen, Natalia de Val, Helen J. Kim, et al. 2013. “A next-Generation Cleaved, Soluble HIV-1 Env Trimer, BG505 SOSIP.664 gp140, Expresses Multiple Epitopes for Broadly Neutralizing but Not Non-Neutralizing Antibodies.” PLoS Pathogens 9 (9): e1003618.

Sanders, Rogier W., Marit J. van Gils, Ronald Derking, Devin Sok, Thomas J. Ketas, Judith A. Burger, Gabriel Ozorowski, et al. 2015. “HIV-1 VACCINES. HIV-1 Neutralizing Antibodies Induced by Native-like Envelope Trimers.” Science 349 (6244): aac4223.

Sanders, Rogier W., and John P. Moore. 2021. “Virus Vaccines: Proteins Prefer Prolines.” Cell Host & Microbe 29 (3): 327–33.

Sliepen, Kwinten, Laura Radić, Joan Capella-Pujol, Yasunori Watanabe, Ian Zon, Ana Chumbe, Wen-Hsin Lee, et al. 2022. “Induction of Cross-Neutralizing Antibodies by a Permuted Hepatitis C Virus Glycoprotein Nanoparticle Vaccine Candidate.” Nature Communications 13 (1): 7271.

Stavenhagen, Kathrin, Hannes Hinneburg, Morten Thaysen-Andersen, Laura Hartmann, Daniel Varón Silva, Jens Fuchser, Stephanie Kaspar, Erdmann Rapp, Peter H. Seeberger, and Daniel Kolarich. 2013. “Quantitative Mapping of Glycoprotein Micro-Heterogeneity and Macro-Heterogeneity: An Evaluation of Mass Spectrometry Signal Strengths Using Synthetic Peptides and Glycopeptides.” Journal of Mass Spectrometry: JMS 48 (6): 627–39.

Struwe, Weston B., Alexandra Stuckmann, Anna-Janina Behrens, Kevin Pagel, and Max Crispin. 2017. “Global N-Glycan Site Occupancy of HIV-1 gp120 by Metabolic Engineering and High-Resolution Intact Mass Spectrometry.” ACS Chemical Biology 12 (2): 357–61.

Sun Weina, Kirkpatrick Ericka, Ermler Megan, Nachbagauer Raffael, Broecker Felix, Krammer Florian, and Palese Peter. 2019. “Development of Influenza B Universal Vaccine Candidates Using the ‘Mosaic’ Hemagglutinin Approach.” Journal of Virology 93 (12): e00333–19.

Thornlow, Dana N., Andrew N. Macintyre, Thomas H. Oguin, Amelia B. Karlsson, Erica L. Stover, Heather E. Lynch, Gregory D. Sempowski, and Aaron G. Schmidt. 2021. “Altering the Immunogenicity of Hemagglutinin Immunogens by Hyperglycosylation and Disulfide Stabilization.” Frontiers in Immunology 12 (October): 737973.

Tokatlian, Talar, Benjamin J. Read, Christopher A. Jones, Daniel W. Kulp, Sergey Menis, Jason Y. H. Chang, Jon M. Steichen, et al. 2019. “Innate Immune Recognition of Glycans Targets HIV Nanoparticle Immunogens to Germinal Centers.” Science 363 (6427): 649–54.

Verkerke, Hans P., James A. Williams, Miklos Guttman, Cassandra A. Simonich, Yu Liang, Modestas Filipavicius, Shiu-Lok Hu, Julie Overbaugh, and Kelly K. Lee. 2016. “Epitope-Independent Purification of Native-Like Envelope Trimers from Diverse HIV-1 Isolates.” Journal of Virology 90 (20): 9471–82.

Walls, Alexandra C., Marcos C. Miranda, Alexandra Schäfer, Minh N. Pham, Allison Greaney, Prabhu S. Arunachalam, Mary-Jane Navarro, et al. 2021. “Elicitation of Broadly Protective Sarbecovirus Immunity by Receptor-Binding Domain Nanoparticle Vaccines.” Cell 184 (21): 5432–47.e16.

Watanabe, Akiko, Kevin R. McCarthy, Masayuki Kuraoka, Aaron G. Schmidt, Yu Adachi, Taishi Onodera, Keisuke Tonouchi, et al. 2019. “Antibodies to a Conserved Influenza Head Interface Epitope Protect by an IgG Subtype-Dependent Mechanism.” Cell 177 (5): 1124–35.e16.

Watson, Michael J., Rick Harkewicz, Edgar A. Hodge, Clint Vorauer, Jonathan Palmer, Kelly K. Lee, and Miklos Guttman. 2021. “Simple Platform for Automating Decoupled LC-MS Analysis of Hydrogen/Deuterium Exchange Samples.” Journal of the American Society for Mass Spectrometry 32 (2): 597–600.

Wei, Chih-Jen, Jeffrey C. Boyington, Kaifan Dai, Katherine V. Houser, Melissa B. Pearce, Wing-Pui Kong, Zhi-Yong Yang, Terrence M. Tumpey, and Gary J. Nabel. 2010. “Cross-Neutralization of 1918 and 2009 Influenza Viruses: Role of Glycans in Viral Evolution and Vaccine Design.” Science Translational Medicine 2 (24): 24ra21.

Weidenbacher, Payton A., and Peter S. Kim. 2019. “Protect, Modify, Deprotect (PMD): A Strategy for Creating Vaccines to Elicit Antibodies Targeting a Specific Epitope.” Proceedings of the National Academy of Sciences 116 (20): 9947–52.

Whittle, James R. R., Adam K. Wheatley, Lan Wu, Daniel Lingwood, Masaru Kanekiyo, Steven S. Ma, Sandeep R. Narpala, et al. 2014. “Flow Cytometry Reveals That H5N1 Vaccination Elicits Cross-Reactive Stem-Directed Antibodies from Multiple Ig Heavy-Chain Lineages.” Journal of Virology 88 (8): 4047–57.

Whittle, James R. R., Ruijun Zhang, Surender Khurana, Lisa R. King, Jody Manischewitz, Hana Golding, Philip R. Dormitzer, et al. 2011. “Broadly Neutralizing Human Antibody That Recognizes the Receptor-Binding Pocket of Influenza Virus Hemagglutinin.” Proceedings of the National Academy of Sciences of the United States of America 108 (34): 14216–21.

Widge, Alicia T., Amelia R. Hofstetter, Katherine V. Houser, Seemal F. Awan, Grace L. Chen, Maria C. Burgos Florez, Nina M. Berkowitz, et al. 2023. “An Influenza Hemagglutinin Stem Nanoparticle Vaccine Induces Cross-Group 1 Neutralizing Antibodies in Healthy Adults.” Science Translational Medicine 15 (692): eade4790.

Wrapp, Daniel, Nianshuang Wang, Kizzmekia S. Corbett, Jory A. Goldsmith, Ching-Lin Hsieh, Olubukola Abiona, Barney S. Graham, and Jason S. McLellan. 2020. “Cryo-EM Structure of the 2019-nCoV Spike in the Prefusion Conformation.” Science 367 (6483): 1260–63.

Wu, Nicholas C., and Ian A. Wilson. 2017. “A Perspective on the Structural and Functional Constraints for Immune Evasion: Insights from Influenza Virus.” Journal of Molecular Biology 429 (17): 2694–2709.

Wu, Nicholas C., Jia Xie, Tianqing Zheng, Corwin M. Nycholat, Geramie Grande, James C. Paulson, Richard A. Lerner, and Ian A. Wilson. 2017. “Diversity of Functionally Permissive Sequences in the Receptor-Binding Site of Influenza Hemagglutinin.” Cell Host & Microbe 21 (6): 742–53.e8.

Yassine, Hadi M., Jeffrey C. Boyington, Patrick M. McTamney, Chih-Jen Wei, Masaru Kanekiyo, Wing-Pui Kong, John R. Gallagher, et al. 2015. “Hemagglutinin-Stem Nanoparticles Generate Heterosubtypic Influenza Protection.” Nature Medicine 21 (9): 1065–70.

Zost, Seth J., Juhye Lee, Megan E. Gumina, Kaela Parkhouse, Carole Henry, Nicholas C. Wu, Chang-Chun D. Lee, et al. 2019. “Identification of Antibodies Targeting the H3N2 Hemagglutinin Receptor Binding Site Following Vaccination of Humans.” Cell Reports 29 (13): 4460–70.e8.

